# Lactate metabolic coupling between the endplates and nucleus pulposus via MCT1 is essential for intervertebral disc health

**DOI:** 10.1101/2025.02.24.640004

**Authors:** Maria Tsingas, Konstantinos Tsingas, Wujuan Zhang, Aaron R. Goldman, Makarand V. Risbud

## Abstract

During skeletal growth, there is an increased secretion of lactate by glycolytic nucleus pulposus (NP) cells of the intervertebral disc. To investigate the role of this anion, we generated annulus fibrosus (AF) and endplate (EP) specific *Mct1^cKO^* (*Slc16a1^Col2CreERT2^*) mice. Histological and spatial transcriptomic studies indicated significant disc degeneration in *Mct1^cKO^*, characterized by NP cell loss and delayed EP maturation. Metabolic assays showed that while AF and EP cells were glycolytic, EP chondrocytes readily metabolized lactate. In EP cells, lactate promoted protein, and H3K18 lactylation, implying epigenetic programming. These findings suggest that NP-derived lactate promotes EP cartilage transdifferentiation into the subchondral bone, and in its absence, continued glucose consumption by the persistent EP cartilage reduces glucose availability to the NP and AF likely contributing to tissue degeneration. This study provides the first *in vivo* evidence that metabolic coupling between NP and EP cells is essential for disc growth and health.

## Introduction

The intervertebral disc, the largest avascular organ in the body, is inherently hypoxic. Glucose, the primary energy source for the disc (1–3), diffuses through the vasculature of the bony endplates into the adjacent endplate cartilage and then into the nucleus pulposus (NP) and annulus fibrosus (AF) compartments. The central NP compartment is primarily cellular at birth, but it accumulates a proteoglycan-rich matrix by skeletal maturity. In tandem, in the AF compartment, cells secrete a collagen-rich matrix and develop into an organized, concentric, lamellar structure that persists throughout life. The embryonic endplate is a homogenous cartilaginous tissue that differentiates into two distinct structures: the cartilaginous endplate (CEP) consisting of 1-2 layers of flattened cells retained throughout life, and transient cartilage consisting of hypertrophic chondrocytes, which transdifferentiate into osteoblasts giving rise to an ossified vascularized bony endplate by skeletal maturity (4–6). During this period of skeletal growth, metabolic demands are likely to be elevated since the disc cells synthesize considerable quantities of extracellular matrix (ECM).

Lactate, the product of anaerobic metabolism, effluxes out of the cell via monocarboxylate transporter 4 (MCT4), a hypoxia-inducible transporter enriched in glycolytic cells (7,8). We have demonstrated the importance of MCT4 in lactate clearance in the NP; inhibition results in metabolic rewiring, lactic acidosis, and subsequent disc degeneration (15). Moreover, *Mct4*-KO mice displayed dysregulated endplate maturation with the persistence of hypertrophic cartilage that fails to differentiate into bone (9). In a recent study, Wang et al. speculated that the lactate from the NP compartment is utilized by the neighboring AF cells, implying a metabolic coupling between compartments (10). However, this scenario is paradoxical considering that the AF, like the NP, is avascular and, thus, unlikely to rely on oxidative metabolism (11). This raises an interesting possibility that NP-derived lactate could be utilized as a metabolic fuel by the bordering endplate cells (9). Indeed, MCT1, a bi-directional transporter enriched in EP and AF preferentially imports lactate across the plasma membrane (10,12).

A recent study showed that as growth plate chondrocytes progress from a proliferative to a hypertrophic state before transdifferentiating into bone, they metabolically rewire from glycolysis to OXPHOS via IGF2 as a bioenergetic switch (13). Another study suggested that Sox9-mediated glutamine metabolism is essential for growth plate chondrocyte function, including gene expression, biosynthesis, and redox homeostasis (14). Relatedly, lactate, which is known to promote glutamine metabolism, can stimulate chondrocyte maturation or act as a metabolic fuel on its own (15). Lactate serves as a metabolic switch in cancer, increasing pyruvate entry into the TCA cycle while also suppressing glycolysis (16). Hui et al. developed a tissue-level flux model in mice, which showed that circulating glucose-derived lactate feeds the TCA cycle in most tissues except for the brain (17). Lactate also serves as a signaling molecule promoting histone and non-histone protein lactylation, via generation of lactyl-CoA groups, which affects gene expression (18). In recent years, studies have begun to address the mechanisms underlying lactylation. For example, lactyl-CoA modification of histones was found to be promoted by SOX9 and YAP/TEAD binding to the enhancer regions of neural crest cells (19), while HBO1 catalyze lysine lactylation on histone H3K9 to regulate gene expression (20). Thus, lactate, which was once considered a waste product, is now considered to serve as the “fulcrum of metabolism” (21).

We hypothesized that the increased abundance of lactate, due to elevated glycolysis in NP, supports EP maturation during skeletal growth. To investigate this, we characterized the spinal phenotype of EP and AF-targeted MCT1 (22) conditional knockout mice, *Slc16a1^Col2CreERT2^* (*Mct1^cKO^*) (23). Histological analyses showed that there was prominent disc degeneration in *Mct1^cKO^* mice characterized by a significant loss of NP cells and an aberrant matrix. Strikingly, the discs of *Mct1^cKO^*mice showed a delay in the formation of bony endplates and the persistence of cartilaginous tissue (13,14,24). These findings provide the first evidence for a growth-linked metabolic coupling between the NP cells and EP chondrocytes. In this way, glucose, spared by endplate cells, can be utilized by cells of the NP and AF, which, in return, generate lactate essential for endplate metabolism and maturation, contributing to the maintenance of disc health.

## Results

### *Mct1^cKO^* mice show pronounced disc degeneration and striking endplate alterations

To determine the fate of lactate and to understand the consequences of restricted lactate uptake, we generated AF- and EP-targeted MCT1 knockout mice. Tamoxifen was injected at 2 weeks postnatally to induce Cre-mediate recombination, and the phenotypes of 6- and 12-month-old *Col2a1^CreERT2^Slc16a1^f/f^*(*Mct1^cKO^*) and *Slc16a1^f/f^* (*Mct1^CTR^*) mice were assessed (Fig. 1a). In a few crosses, we used an Ai9 tdTomato+ reporter to mark Cre-recombinase activity. As expected, tdTOM+ cells were robustly present in the AF and EP in control (*Ai9^tdTomato/+^Col2a1^CreERT2^*) and knockout discs. To confirm the protein deletion, we stained for MCT1 and observed an appreciable loss of staining in the AF and EP cells of *Mct1^cKO^* discs (Fig. 1b). Modified Thompson Grading of Safranin O/Fast Green stained *Mct1^cKO^* caudal discs showed pronounced NP degeneration and loss of demarcation between the NP and AF compartments. At both 6- and 12-months, *Mct1^cKO^*endplates displayed pronounced Safranin-O-positive staining in the bony endplate region, indicating the persistence of cartilaginous tissue. NP and AF had higher grades of degeneration, by both distribution and average, along with a strikingly higher number of discs showing the presence of cartilaginous remnants in the EP of *Mct1^cKO^* (Fig. 1c-e). Moreover, in *Mct1^cKO^* mice, cartilaginous tissue occupied a significantly higher area of the EP and showed higher grades of EP degeneration (Fig. 1f). Finally, TUNEL staining revealed a higher incidence of cell death in the NP and EP of *Mct1^cKO^* mice (Fig. 1g). In summary, loss of MCT1 significantly impacted cellular and extracellular constituents of the intervertebral disc.

**Figure 1.**
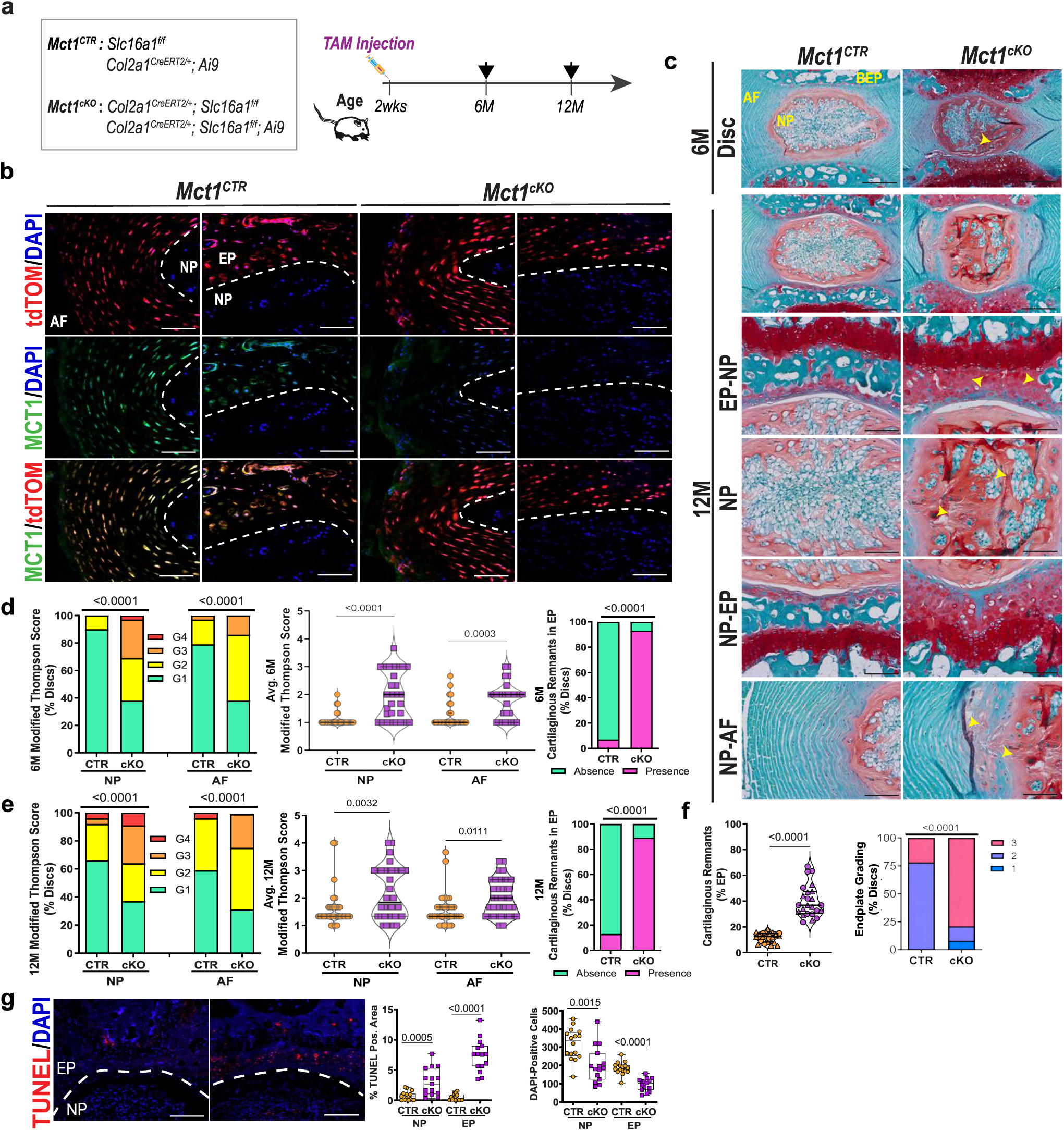
*Mct1^cKO^* exhibit pronounced disc degeneration and striking endplate alterations. a) Overview of the timeline for the generation of *Mct1^CTR^*(CTR) and *Mct1^cKO^* (cKO) mice. b) Colocalization of Ai9 (tdTOM+) reporter marking Cre recombinase activity with MCT1 shows MCT deletion in the AF and EP compartments of cKO mice (scale bar = 50 μm). c) Safranin O/Fast Green staining of the discs from 6- and 12- month-old *Mct1^CTR^* and *Mct1^cKO^* mice. Yellow arrowheads indicate degenerative changes. Scale bar: Rows 1 and 2 = 100 μm; Rows 3-6 = 50 μm. d, e) Distribution and average modified Thompson grades, and incidence of EP phenotype in (d) 6-month-old and (e) 12-month-old mice. N; 6M: 9 mice/genotype (5 males, 4 females), 3-4 caudal discs/mouse, n;12M= 11-12 mice/genotype (CTR: 6 males, 5 females; cKO: 6 males, 6 females), 2-4 caudal discs/animal were assessed. f) Area of persisting EP cartilage and cartilage grading at 12-month n = 11-12 mice/genotype. g) Representative TUNEL staining images showing apoptotic cells in the NP and EP regions of disc sections and total number of cells in NP and EP compartments (scale bar = 50 μm), *n* = 5 mice/genotype (3 males, 2 females), 3-4 discs/animal. Data represented as violin plots with median and quartiles. Significance for the grading distribution was determined using Fisher’s Exact test, all other comparisons were performed using an unpaired t-test or Mann-Whitney test based on data normality.

### MCT1 loss delays endochondral ossification and formation of the bony EP

To characterize the phenotype, immunohistological analyses were performed on 12-month-old *Mct1^cKO^* mice. Notably, EP showed persistence of hypertrophic cartilage as evidenced by robust COL X expression alongside decreased EMCN and TNAP staining, suggesting fewer blood vessels, retention of cartilaginous phenotype and decreased bone formation (Fig. 2a). Consistent with significant NP cell loss and altered morphology, the abundance of NP phenotypic marker GLUT1 was markedly reduced in the *Mct1^cKO^* discs. Further, the staining of lactate efflux channel MCT4 and enzymes involved in lactate metabolism, LDHA and LDHB, were reduced in the *Mct1^cKO^* discs (Fig. 2b). These findings suggest that the inhibition of lactate uptake through MCT1 is sufficient to drive metabolic changes that may contribute to the delayed endochondral ossification observed in the endplate.

**Figure 2.**
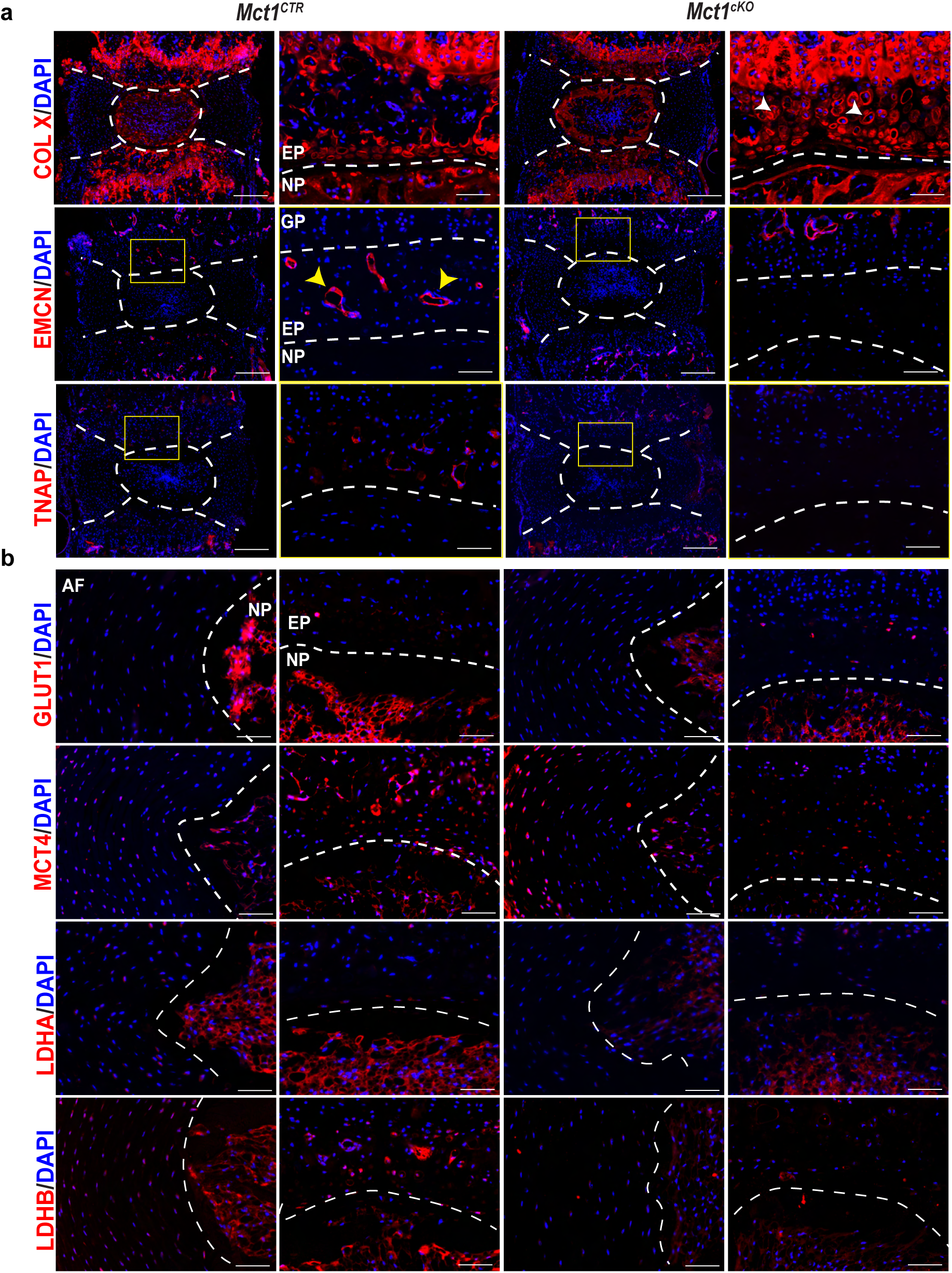
MCT1 loss delays endochondral ossification and formation of the bony EP. a) Representative images of COL X, EMCN, and TNAP in the endplate region of 12-month-old *Mct1^CTR^*and *Mct1^cKO^* mice. White arrowheads indicate hypertrophic chondrocytes, yellow arrowheads show blood vessels. Scale bar: columns 1, 3 = 200 μm, columns 2, 4, = 50 μm. b) Representative images showing GLUT1, MCT4, LDHA, and LDHB in the AF and NP compartments, scale bar = 50 μm. *n* = 4 mice/genotype (2 males, 2 females), 3 discs/animal assessed. Dotted lines indicate boundaries between different tissue compartments within the disc.

### *Mct1^cKO^* discs show disorganized ECM and NP fibrosis

At 12 months, *Mct1^cKO^* discs showed significant structural changes, including an increase in thin collagen fibrils, a reduction in intermediate fibril thickness, and notable collagen fibril disorganization in the AF (Fig. 3a-b). There was also a marked increase in the incidence and extent of fibrosis of the NP accompanied by increased COL I expression (Fig. 3a-c). To determine the quality of the collagen matrix, we used a fluorescently labeled peptide (CHP) that binds to denatured collagen (25). In the EP, higher CHP binding indicated slight disorganization and an increased abundance of denatured collagen in *Mct1^cKO^* mice (Fig. 3c). Furthermore, marked alterations in aggrecan and chondroitin sulfate staining patterns within NP of *Mct1^cKO^* mice highlighted matrix disorganization (Fig. 3d).

**Figure 3.**
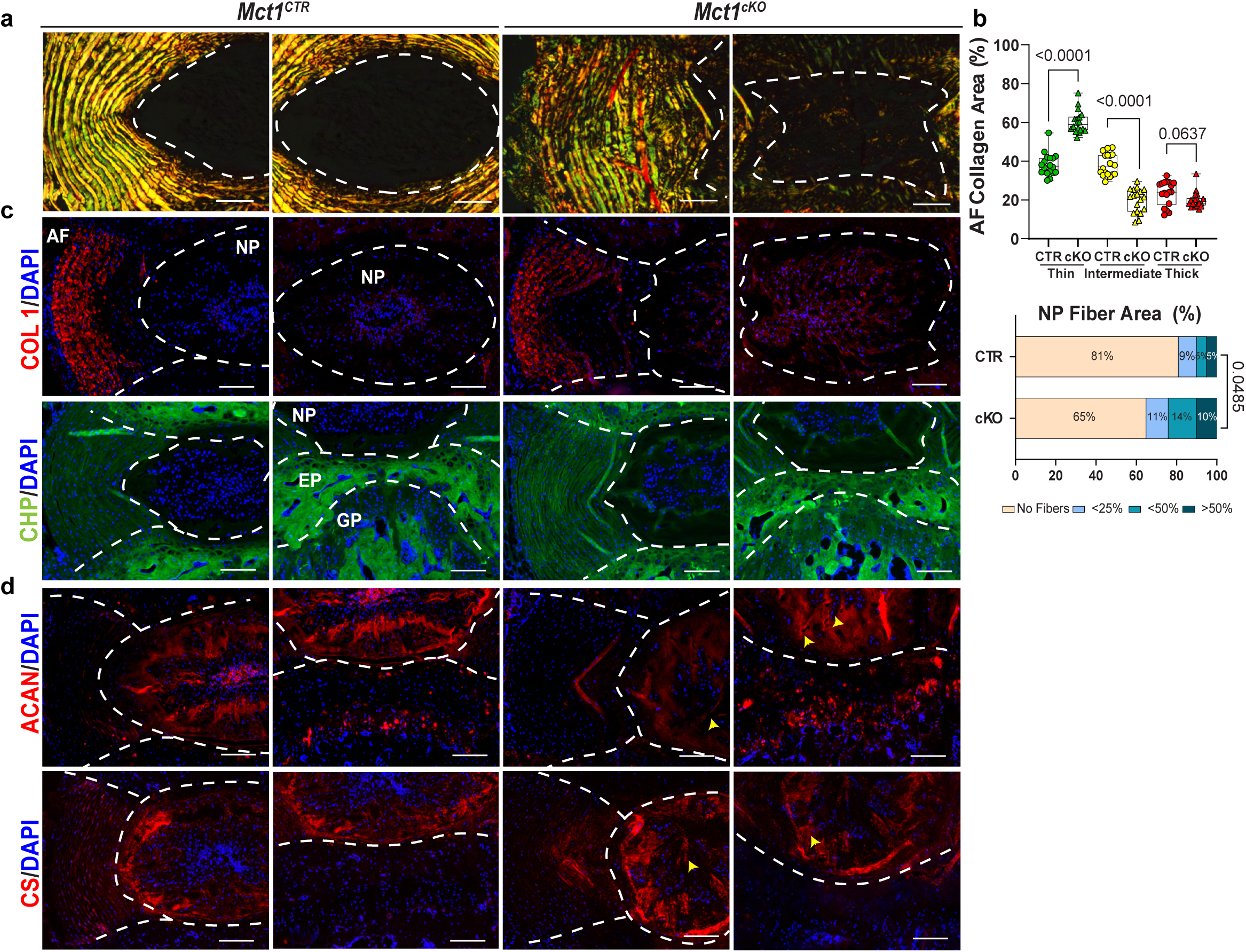
Mct1^cKO^ discs exhibit disorganized matrix and NP fibrosis. a) Polarized imaging of picrosirius red stained discs of 12-month-old *Mct1^CTR^* and *Mct1^cKO^*mice showing collagen organization, scale bar = 100 μm. b) Quantification of AF fiber thickness distribution and the NP fibrosis incidence. c) Representative images of 12-month-old discs showing collagen I (COL I), collagen hybridizing peptide (CHP), aggrecan (ACAN), chondroitin sulfate (CS) staining, scale bar = 100 μm, *n* = 3 mice (2m, 1f)/genotype, 2-3 discs/mouse. Yellow arrowheads indicate degenerative changes. Data represented as violin plot with median and quartiles (b). Significance for AF collagen area was determined by using an unpaired t-test or Mann-Whitney, as appropriate. Significance for NP fiber area distribution was determined using a χ^2^ test. Dotted lines mark the boundaries between the different tissue compartments within the disc.

### MCT1 loss affects endplate mineralization but does not markedly impact vertebral bone

To evaluate whether the endplate phenotype is restricted to the subchondral region or extends into vertebral bone, μCT analysis was performed on 12-month-old mice. Reconstructed three-dimensional scans revealed negligible changes in vertebral bone morphology of *Mct1^cKO^*mice (Supplementary Fig. 1a). There were no changes in disc height or disc height index (DHI); however, there was a small decrease in vertebral length (Supplementary Fig. 1b). Cortical and trabecular bone properties were assessed and showed no major differences in these parameters, except for significant increases in trabecular and cortical BMD (Supplementary Fig. 1c). *Mct1^cKO^*mice displayed significant decrease in caudal endplate BMD, consistent with the histological outcomes showing the persistence of cartilage (Supplementary Fig. 1d).

### Spatial transcriptomics analysis reveals altered transcriptomic profiles and cell fate in ***Mct1^cKO^ discs*.**

We used spatial transcriptomics to gain further insights into gene expression changes in *Mct1^cKO^* discs (Fig. 4a). This approach allowed for the capture of positional gene information, including crosstalk between disc compartments, which is otherwise lost in scRNA-sequencing techniques. H&E images, with corresponding spatial maps of the *Mct1^CTR^* and *Mct1^cKO^* motion segments, indicated distinct tissue-specific clustering (Fig. 4b) in the region defined between superior and inferior growth plates. Seurat clustering analysis showed 11 distinct cell clusters (Fig. 4c). Clusters were annotated from a merged dataset based on top marker genes (Fig. 4d-e). We observed two distinct NP populations, periNP (pNP) and centriNP (cNP), which clustered distinctly from the rest of the cell types. Chondrocyte, muscle, and osteoblast cell populations were segregated into Clusters “I”, “II”, or “III” and AF and tendon cells presented as single clusters. *Mct1^cKO^* samples showed negligible differences in the clustering of AF cells, and the lack of a tendon cluster was likely due to sample processing. Interestingly, *Mct1^cKO^* showed differences in the positioning of certain clusters in spatial maps, in particular, osteoblasts were largely absent from the bony endplate region of mutant mice, consistent with the histological observation that the bony endplate was poorly structured (Fig. 4b). Further comparisons were performed between *Mct1^CTR^* vs *Mct1^cKO^* genes on validated cluster annotations using bonafide marker genes. These studies highlighted distinct differences between genotypes (Fig. 4f; Supplementary Fig. 2).

**Figure 4.**
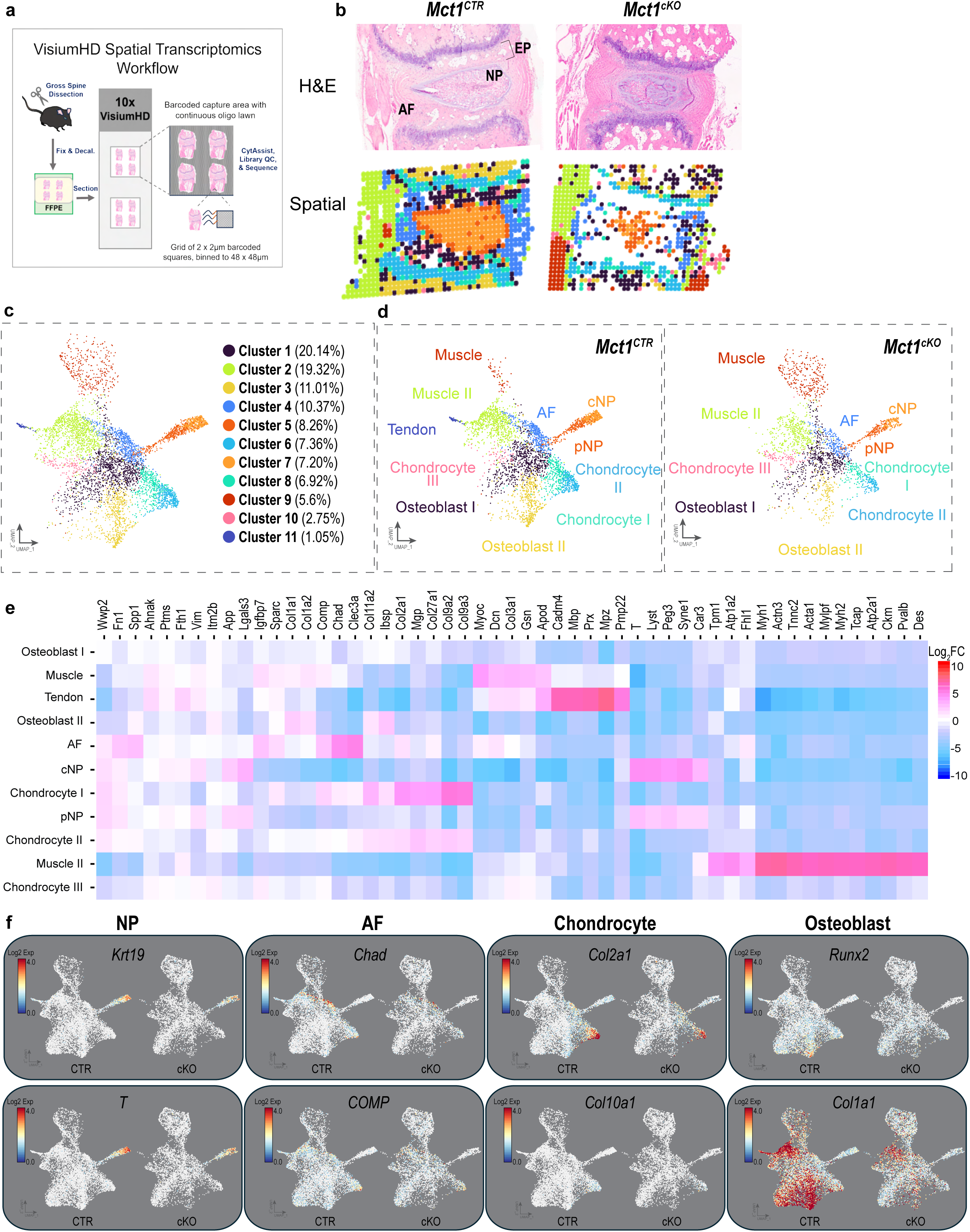
Spatial transcriptomics unveils altered transcriptomic profiles and cell fate in *Mct1^cKO^*. a) Spatial transcriptomics workflow using VisiumHD platform for spinal motion segments of 6-month-old *Mct1^CTR^* and *Mct1^cKO^*mice. b) H&E-stained images of a control and knockout mouse spinal motion segment (Caudal level 7/8) and the corresponding spatial profiles following clustering analysis. c) UMAP showing 11 distinct cell clusters identified by Seurat package using merged data set from CTR and cKO mice. *n* = 4 mice/genotype; 2 m, 2 f. d) Cluster annotated UMAP plots of *Mct1^CTR^* and *Mct1^cKO^* showing distinct cell populations with the disc compartment. e) Heatmap of expressed markers used for cluster annotation. f) Feature plots showing spatial expression of marker genes in the NP, AF, chondrocyte, and osteoblast populations in CTR and cKO mice.

To examine the genotype-based differences between cell clusters with the same annotation, significantly differentially expressed genes (DEGs) were identified (FDR < 0.05). Notably, the ‘cNP’ cluster had 103 downregulated and 20 upregulated DEGs (Supplementary Fig. 3a). Pathway analysis of the cluster DEGs was carried out using the AI CompBio tool (PercayAI Inc., St. Louis, MO) to explore thematic associations between these genes (Fig. 5a-b). A ball-and-stick plot of DEGs from the ‘cNP’ cluster revealed a prominent thematic supercluster, associated with muscle structure and cytoskeletal processes. Key themes within this supercluster included *Degenerative Disc Disease*, *Temporomandibular Joint Articular Cartilage Development*, *Actin Bundling Activity*, and *Fibronectin Fibril Organization*. The top genes contributing to these themes include *Acan*, *Fth1*, *Sdc4*, and *Ibsp*, which highlight significant changes in disc homeostasis. Another distinct supercluster focused on actin and protein binding, featuring themes such as: *Amino-terminal Vacuolar Sorting Propeptide Binding and Actin Capping Protein of Dynactin Complex.* Additionally, a smaller supercluster related to endocytosis and vesicular trafficking encompassed themes like *Acidification of Endocytotic Vesicles and COPI Coating of Golgi Vesicle cis Golgi to rough ER*, with key genes including *Golgb1*, *Cavin1,* and *Cltb.* Another theme *Entrainment of Circadian Clock by Photoperiod* driven by significant changes in *Per1* and *Dbp*, suggested disruptions in circadian rhythm. Overall, these findings indicate that the cNP of *Mct1^cKO^* mice exhibits significant alterations in the cellular phenotype, accompanied by degenerative changes. Gene set enrichment analysis (GSEA) was conducted on DEGs defined by q-values of 0.25 and 0.05. In cNP cells of *Mct1^cKO^* mice, GSEA showed upregulation in pathways related to muscle: ‘actin-myosin and muscle filament sliding’ and ‘muscle contraction’, supporting altered cell phenotype and tissue fibrosis (Supplementary Fig. 3b-c).

**Figure 5.**
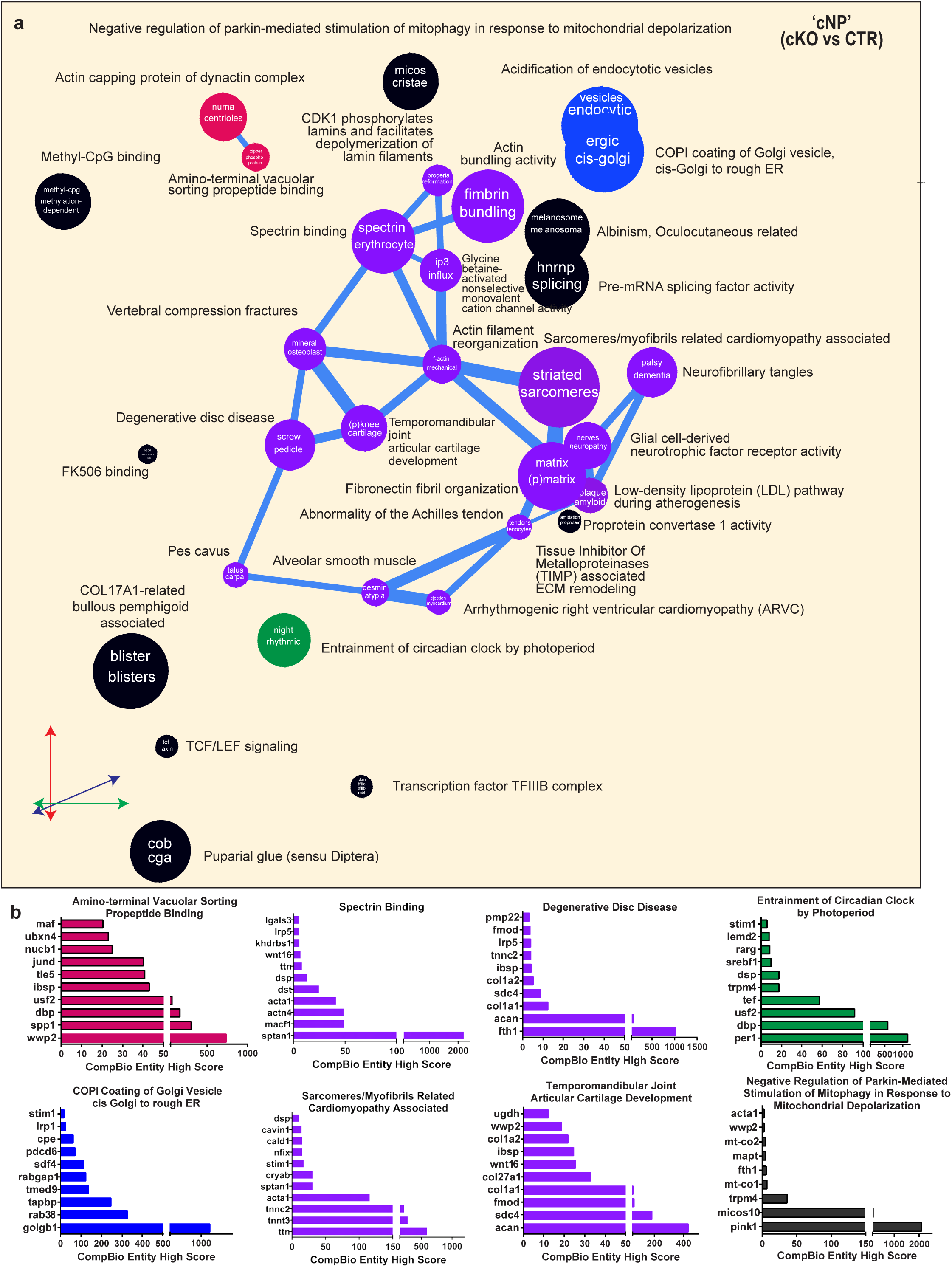
MCT1 deletion dysregulates the transcriptomic signatures associated with ‘central NP’ cell cluster. a) Thematic organization of concepts generated by CompBio, derived from literature analysis of differentially expressed genes in the ‘central NP’ cluster of the *Mct1^cKO^* mouse discs, reveals key physiological processes affected by impaired lactate uptake. Superclusters of themes associated with these processes include homeostasis and development (purple), actin and protein binding (red), endocytosis and vesicular trafficking (blue), and circadian rhythm regulation (green). b) Top DEGs enriched in key themes are displayed as entities based on their CompBio entity high score, 4 mice/genotype; 2 m, 2 f.

We analyzed the differences in the ‘Chondrocyte I’ cluster between genotypes and found 39 downregulated and 3 upregulated DEGs (Supplementary Fig. 3d). The ball-and-stick plot of the ‘Chondrocyte I’ DEGs revealed strong centrality and connectivity with a prominent supercluster including several themes: *Sarcomeres/Myofibrils Related*, *Amyloid-Beta Precursor Protein*, *BMP Signaling*, *Vertebral Compression Fractures,* and *Epiphyseal Dysplasia* (Supplementary Fig. 5a-b). A small ribosome RNA-related supercluster comprising two themes *Cleavage in ITS2 between 5.8S RNA- and Mitochondrial lrRNA Export from Mitochondrion* with high enrichment score seen in *Mt-Atp8*, *Nxf1*, and *Nop53.* Other significantly altered genes, such as *App, Sost, Col1a1, Col1a2, Fgfr3,* and *Ttn*, indicated disruptions in cartilage biology, particularly in chondrocyte development and skeletal dysplasia. Collectively, these findings highlight a distinctive chondrocyte-like phenotype that is marked by alterations in growth factor signaling, endocytosis, and muscle-related pathways. In Chondrocyte I cells of *Mct1^cKO^* mice, GSEA analysis showed increases in ‘positive regulation of gene expression’, ‘muscle contraction’, and ‘skeletal myofibril assembly’ (Supplementary Fig. 3e-f).

### EP chondrocytes utilize lactate as a metabolic fuel in TCA cycle

To better understand the metabolic changes observed *in vivo*, Seahorse assays were performed to measure the rate of glycolytic and oxidative metabolism using primary EP chondrocytes cultured with varying amounts of lactate and under osteogenic differentiation conditions for one week at physioxia (5% O_2_) (26). Results showed that lactate supplementation was not sufficient to drive any major bioenergetic changes except for increased glycolysis (Fig. 6a-b) and increased rate of glycolytic ATP production in the 5 mM lactate group but not at a higher lactate concentration of 10 mM (Fig. 6c). Interestingly, this difference was lost with oligomycin treatment, suggesting the involvement of lactate-dependent mitochondrial activity.

**Figure 6.**
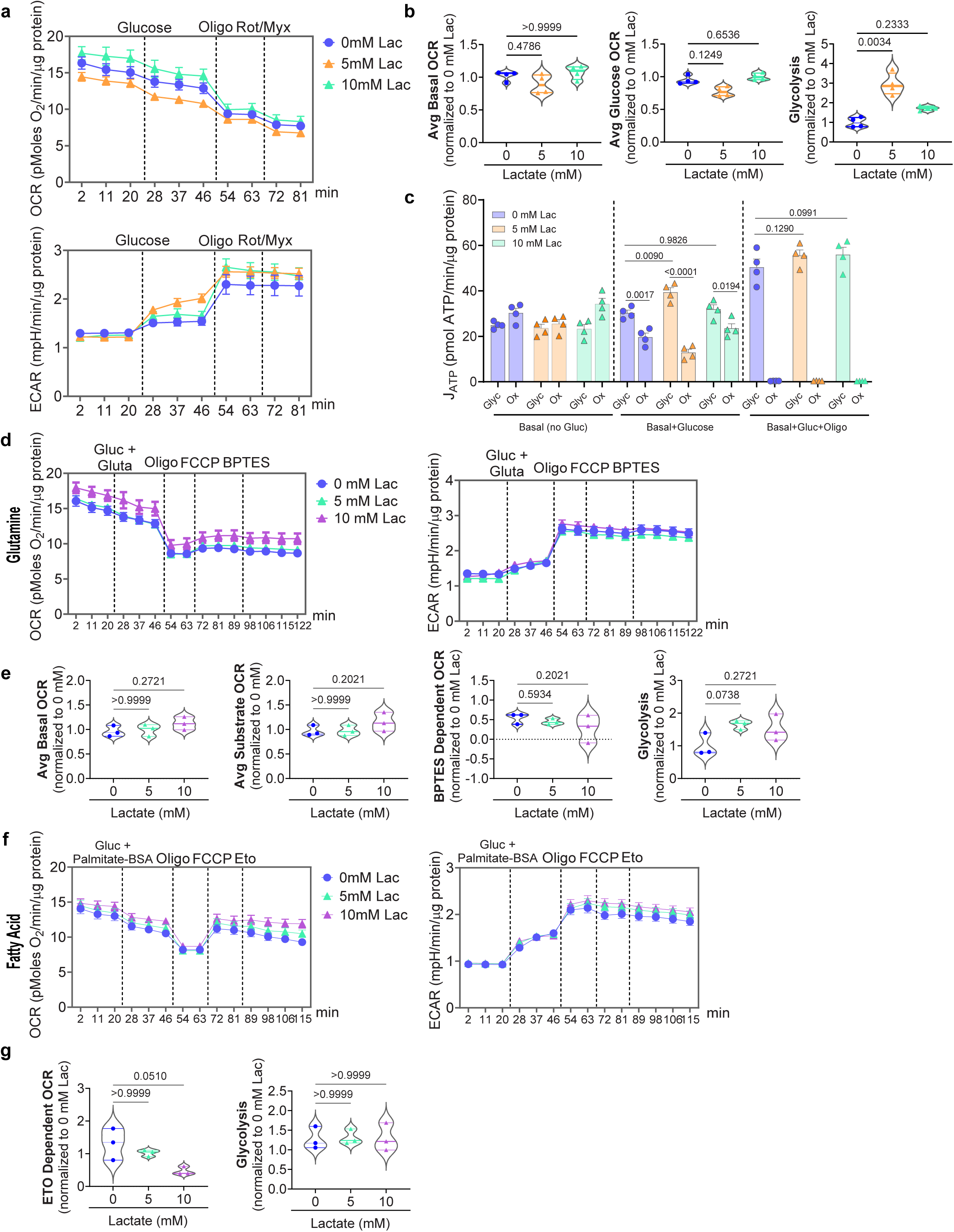
Lactate does not promote OXPHOS switch in EP cells. Primary EP cells were cultured with sodium L-lactate for one week under osteogenic conditions to measure effect on bioenergetics. a) Normalized ECAR and OCR traces, with corresponding (b) Average basal and glucose-dependent OCR, as well as glycolysis. c) ATP production rate partitioned into glycolytic and oxidative ATP under basal, substrate, ATP-linked conditions. d) Normalized OCR and ECAR traces from substrate utilization experiments in a mixed-glucose-glutamine environment, with corresponding (e) average basal and glucose-dependent OCR, BPTES-dependent responses, and glycolysis relative to no-lactate control. f) Normalized ECAR and OCR traces of fatty acid utilization experiment, with corresponding (g) quantification showing etomoxir-dependent OCR and glycolysis parameters. Quantitative measurements are expressed as mean ± SEM (*n* = 4 experiments, with 4 technical repeats/group/trace) (a, c, d, f) and as violin plots with medians and interquartile ranges (b,e,g). Significance was assessed using Kruskal-Wallis test with Dunn’s post hoc analysis (b, e, g) and one-way ANOVA with Sidak’s post hoc test (c).

To assess whether lactate promotes the utilization of glutamine or fatty acids in EP cells, substrate utilization was evaluated. There were no differences in OCR or ECAR traces, BPTES-dependent OCR, and glycolysis (Fig. 6d-e). Similarly, lactate did not appear to interfere with fatty acid utilization as confirmed by etomoxir (ETO) treatment (Fig. 6f-g). Taken together, these experiments suggest that lactate on its own or in the presence of glutamine or fatty acids does not alter common substrate utilization.

Lactate is converted to pyruvate by LDHB activity, which enters the TCA cycle and increases oxidative phosphorylation (OXPHOS). Although both glucose and lactate can generate pyruvate, their relative contributions to EP chondrocyte metabolism are unknown. To determine the fate of lactate, we cultured primary EP cells with [U-^13^C] glucose (5 mM) and increasing concentrations of [3-^13^C] lactate for 30 minutes to measure flux into M+1 and M+3 pyruvate and M+1 and M+2 citrate pools (Fig. 7a). We observed that approximately 50-60% of the pyruvate pool was derived from glucose (M+3), while lactate contributed about 15-20% of pyruvate (M+1) (Fig. 7b). Interestingly, glucose (M+2) and lactate (M+1) contributed about equally (∼8-12%) to the citrate pool (Fig. 7b). These results suggest that in EP cells, only a fraction of glucose-derived pyruvate contributes to the citrate pool via the TCA cycle, whereas a significantly higher proportion of lactate-derived pyruvate enters into the TCA cycle. To assess lactate contribution to the metabolite pools at steady state, we performed a similar experiment where EP cells were cultured with 10 mM [3-^13^C] lactate and 5 mM unlabeled glucose for 24 hours (Fig. 7c). We observed that there was further enrichment of labeling into pyruvate and citrate pools, and lactate also contributed to aspartate and malate pools (Fig. 7d), suggesting that lactate contributes as a carbon source to the intermediates of the TCA cycle. Together, these experiments suggest that EP cells utilize both glucose and lactate as substrates, highlighting their mixed glycolytic and oxidative metabolism.

**Figure 7.**
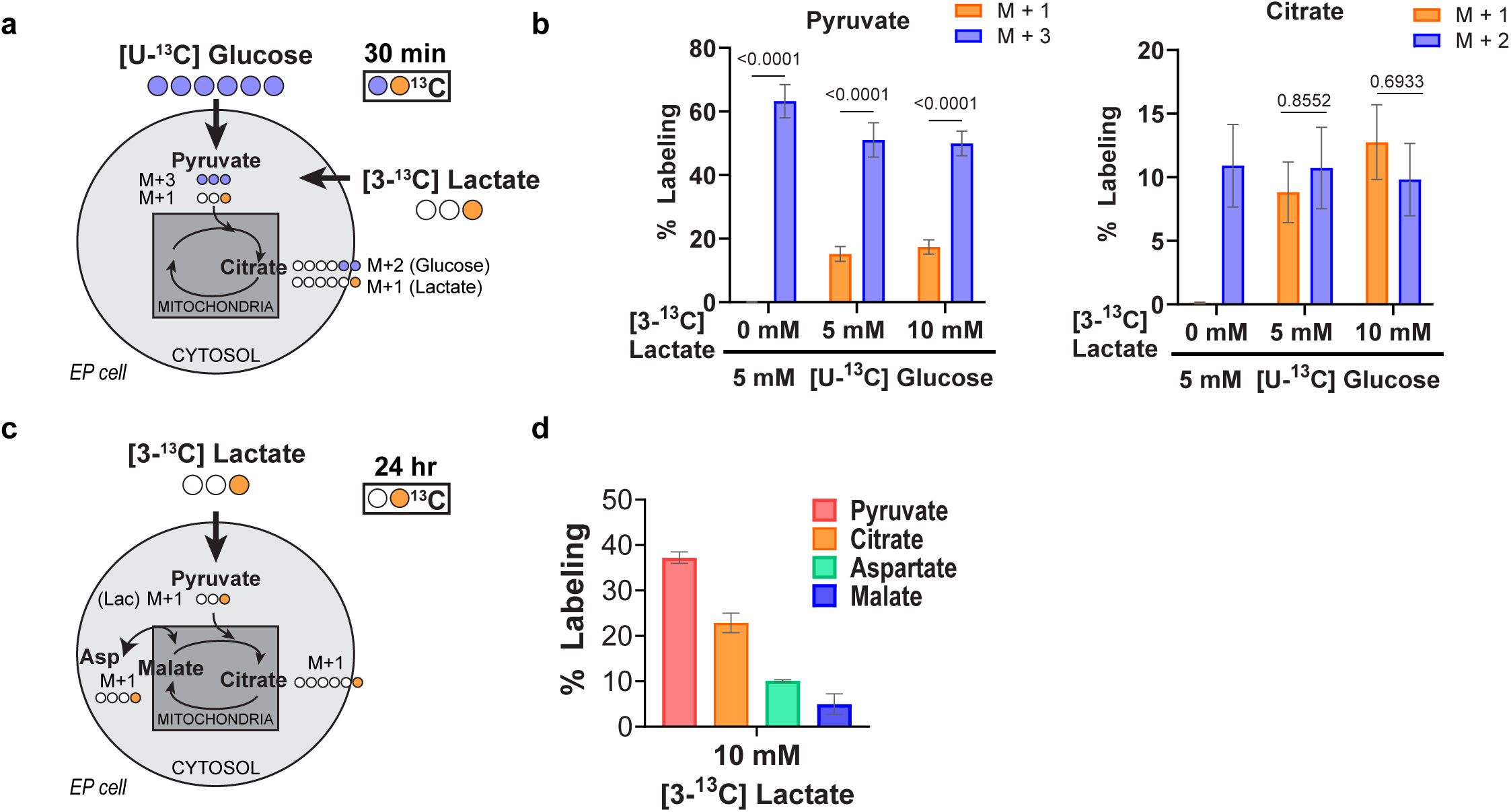
EP cells use lactate in TCA cycle. a) A schematic diagram of the dual [U-^13^C] glucose and [3-^13^C] lactate tracing, demonstrating the flow of carbons into the TCA cycle. b) Percent labeling of citrate and pyruvate after 30 min from 5 mM [U-^13^C] glucose with increasing concentrations of [3-^13^C] lactate (up to 10 mM) in EP cells. Significance was determined using One-way ANOVA with Sidak’s post hoc test for multiple comparisons. c) Schematic diagram showing [3-^13^C] lactate tracing, illustrating carbon flow into the TCA cycle. d) Percent labeling into TCA metabolites after 24 hr from 10 mM [3-^13^C] glucose in presence of 5 mM unlabeled glucose in EP cells. Data are presented as the mean ± SEM, *n* = 5 independent samples/condition.

### AF cells are glycolytic and do not prefer lactate as a metabolic substrate

We performed identical double labeling experiments with [U-^13^C] glucose and [3-^13^C] lactate under physioxic conditions (5% O_2_) to examine if AF cells utilize lactate (Supplementary Fig. 5a). We observed that glucose was the major contributor to pyruvate (M+3, ∼66%), lactate (M+3, ∼55%), and citrate (M+2, ∼15-20%) pools. Only ∼21% of the intracellular lactate was derived from tracer uptake (M+1), and it contributed minimally to both pyruvate (M+1, ∼2-8%) and citrate (M+1, ∼1-5%) pools, suggesting that AF cells are predominantly glycolytic and prefer glucose as their primary carbon source (Supplementary Fig. 5b).

We then assessed the effects of lactate supplementation on AF cell glycolysis and bioenergetics using Seahorse assays, cultured with varying amounts of lactate for 24 hours. Upon the introduction of glucose, lactate-supplemented cells showed a prominent increase in ECAR, signifying increased glycolysis due to a preference for glucose. However, this difference was lost when mitochondria were inhibited with oligomycin and rotenone/myxothiazol; thus, the glycolytic and reserve capacities or any OCR-related parameters were not affected (Supplementary Fig. 5c, d). When cellular bioenergetics were assessed, the proportions of glycolytic to oxidative ATP in the lactate-supplemented groups, following the glucose bolus, evidenced an increased proportion of glycolytic versus oxidative ATP relative to the control group (Supplementary Fig. 5e). This suggested that the continued presence of lactate increased cellular glycolysis and, thus, increased the rate of glycolytic ATP production, as soon as glucose became available. As a compensatory response to loss of ATP through OXPHOS, the rate of glycolytic ATP production increased in all groups when mitochondrial ATP synthase was inhibited with oligomycin, and the differences between lactate and control groups were no longer evident.

### Lactate promotes protein and histone lactylation in EP chondrocytes and alters their transcriptomic program

To investigate if lactate serves as a signaling molecule, we assessed if exogenous lactate promoted protein and histone lactylation, specifically at H3K18 (Fig. 8a). Lactate supplementation induced a clear increase in pan-lactylation with a negligible difference in pan-acetylation (Fig. 8b-d). A similar pattern was observed with histone modifications, where there was a notable increase in lactylated levels of H3K18, with no observable differences in H3K27 (Fig. 8e-f). Immunofluorescence staining showed prominent nuclear staining of H3K18la in lactate-treated cells, whereas H3K27ac showed no discernible differences (Fig. 8g-f). Noteworthy, we observed abundant pan-Kla staining and nuclear H3K18la localization in all disc compartments of 6-month-old *Mct1^CTR^*mice, with a marked reduction in the *Mct1^cKO^* mice (Fig. 8i), suggesting that lactate promotes protein and histone lactylation of disc cells *in vivo*.

**Figure 8.**
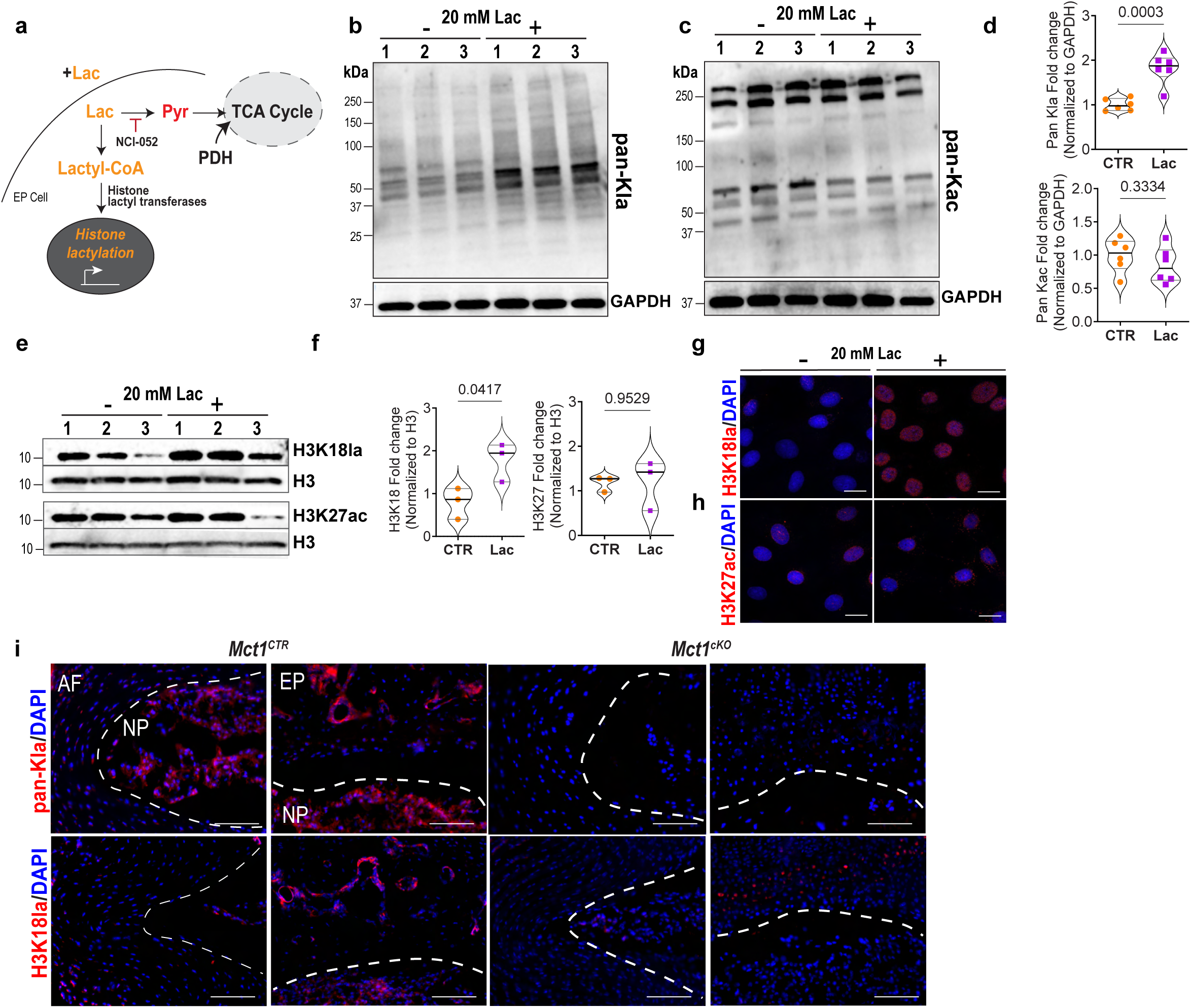
Lactate treatment in endplate cells induces lactylation. (a) A schematic diagram of lactylation via lactate-derived lactyl-CoA within an endplate cell. Lactate contributes to the formation of lactyl-CoA, which can be transferred to histones, by histones lactyl transferases, leading to histone lactylation, affecting gene expression. (b-c) Western blot of pan-lactylation (pan-Kla) and pan-acetylation (pan-Kac) in total protein from endplate cells treated with 20 mM sodium L-lactate or NaCl for 24 hours in physioxia. (d) Densitometric analysis of protein fold changes normalized to GAPDH, *n* = 6 independent samples/condition. (e-f) Western blot and densitometric analysis of histones extracted from lactate-treated endplate cells, probed for H3K18la and H3K27ac, *n* = 3 independent samples/condition. (g-h) IF staining of H3K18la and H3K27ac in endplate cells treated with 20 mM sodium L-lactate or NaCl for 24 hours in physioxia, scale bar = 15 μm. (i) Representative images of 6-month-old discs showing pan-lactylation (pan-Kla) and H3K18la, nuclear-specific for lactylation, in the AF-NP and EP, scale bar = 50 μm, *n* = 3 mice/genotype (2m, 2f), 2-3 discs/mouse. Quantitative measurements are shown as violin plots with medians and interquartile ranges.

To explore the effects of lactate supplementation on the transcriptomic landscape, we performed bulk RNA-Seq on endplate chondrocytes treated with 20 mM lactate for 24 hours. There were 131 DEGs (FDR < 0.05), 80 downregulated and 51 upregulated. Hierarchical clustering showed that lactate-treated and untreated groups clustered distinctly (Supplementary Fig. 6a-b). We then investigated the level of expression of genes encoding key enzymes involved in lactylation, including a recently described lactyl-CoA synthetase ACSS2 and several known lactyl transferases. The heatmap of TPM values shows a robust expression of *Acss2* by EP cells and a clear reduction in lactate-treated cells. EP cells expressed several lysine acetyl-transferases (KAT) that also serve as lactyl-transferases, including *Kat2a*, *Kat7* and *Kat5,* with *Kat5* showing the highest expression (Supplementary Fig. 6c). We analyzed the biological context of the upregulated and downregulated DEGs using the CompBio tool. Interestingly, analysis of the downregulated DEGs showed strong enrichment around *Very-long-chain Fatty Acyl-CoA oxidase activity*, *Fatty Acid Beta-Oxidation*, *Peroxisome Proliferator-Activated Receptor (PPAR) agonists*, *Insulin-like Growth Factor (IGF) Activity Regulation by IGFBPs*, *LPA/Lysophospholipid Signaling,* and *All-trans Retinoic Acid 18-hydroxylase Activity* (Fig. 9a). Strong gene signals from bone matrix protein, *Spp1*, were shown to be downregulated along with *Ankh*, suggesting decreased extracellular pyrophosphate. Other top genes within these thematic clusters include *Enpp2, Fads2*, *Acta1*, *Abca1*, *Scd*, *Igfbp5*, *Srebf1*, *Dhrs3 (All-trans Retinoic Acid 18-hydroxylase Activity),* and *Mgll (Digestion of Cholesterol Esters)* (Fig. 9b). Key findings highlighted the roles of fatty acids, retinoic acid, pyrophosphate, calcium-dependent enzyme activities, and tissue development. Overall, the gene expression profile indicated lactate-mediated downregulation of fatty acid-related genes and suggested that the cartilage and sclerotomal phenotype was suppressed in endplate cells.

**Figure 9.**
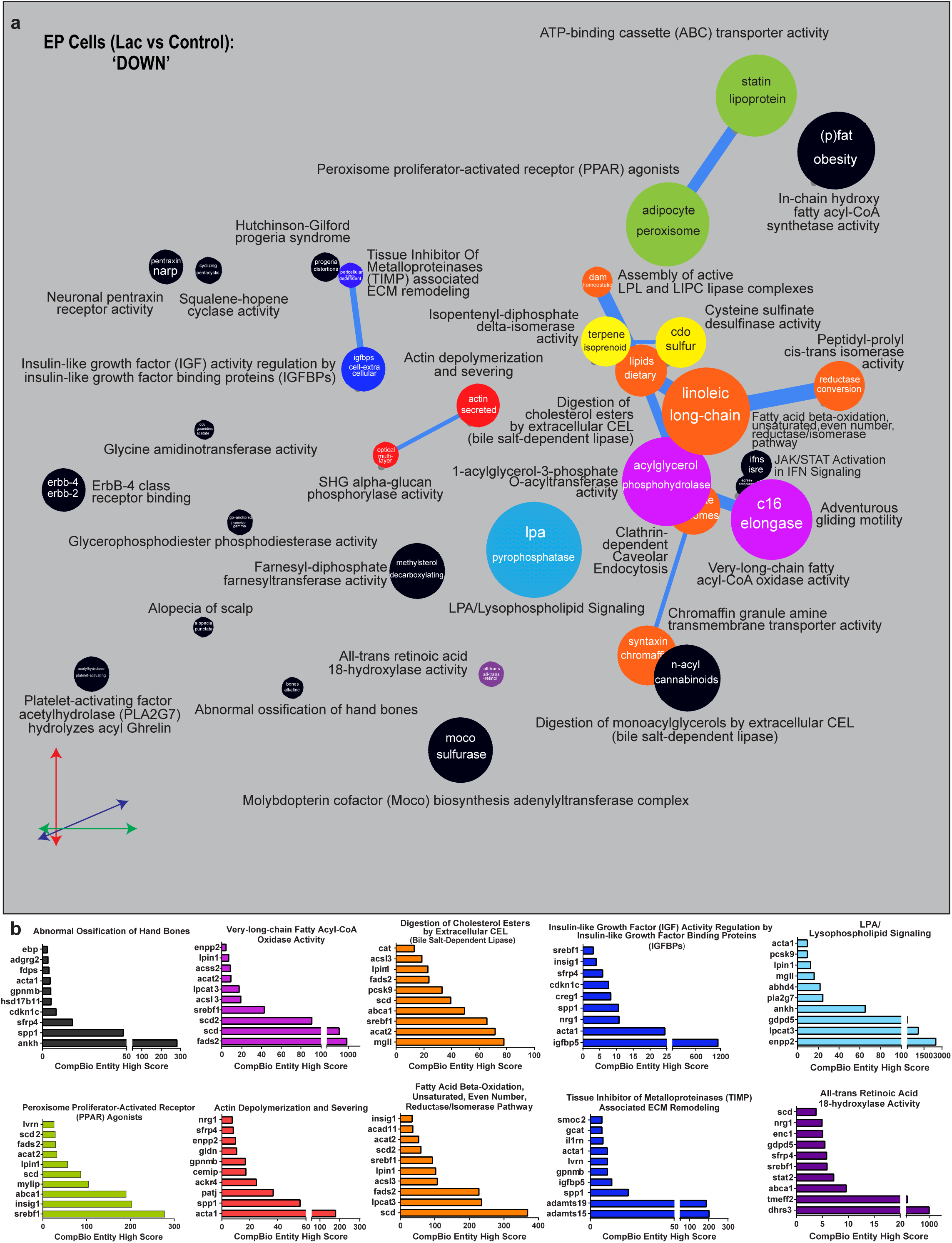
Lactate supplementation suppresses the cartilage sclerotomal phenotype in EP cells. (a) CompBio-generated thematic organization of concepts, derived from literature analysis of differentially downregulated genes in lactate-treated EP cells, reveals key physiological processes affected by lactate. Superclusters of themes associated with these processes include: fatty acids (orange), metabolic enzymes (magenta), calcium-dependent enzyme activities (yellow) ATP-binding cassette transporter activity (green), and growth and tissue remodeling regulation (blue), structural assembly (red). retinoic acid (purple), pyrophosphate (light blue). (b) Top DEGs enriched in key themes are displayed as entities based on their CompBio entity high score, (*n* = 5 experiments; 2 conditions: 20 mM lactate or NaCl).

While upregulated DEGs showed slightly less pronounced thematic enrichment, many converged on pathways related to *Growth Factor Activity, Establishment of Mitotic Spindle Orientation Involved in Growth Plate Cartilage Chondrocyte Division, Axial Skeleton Plus Cranial Skeleton, Follistatin/Inhibin/Activin Axis, Mesenchymal Stem Cell Maintenance Involved in Nephron Morphogenesis* (Supplementary Fig. 7a). Central ideas were focused on immune signaling pathways, inflammatory responses, and growth factor regulation, emphasizing molecular mechanisms that drive cell migration, activation, and replication inhibition. Significant changes in *Fgfr1*, *Csf1*, *Ccl2, Col2a1*, and *Meox1* suggested significant alterations in growth regulation and development (Supplementary Fig. 7b). These findings revealed changes in the EP cell phenotype induced by lactate treatment.

In summary, our study, for the first time, highlights an MCT-dependent metabolic coupling between the EP, NP, and AF, wherein glucose and lactate serve as the primary metabolic currency. Consequently, in *Mct1*^cKO^ mice, the dysregulation of lactate coupling leads to impaired bony endplate formation, pronounced NP cell loss, and disc degeneration (Fig. 10a).

**Figure 10.**
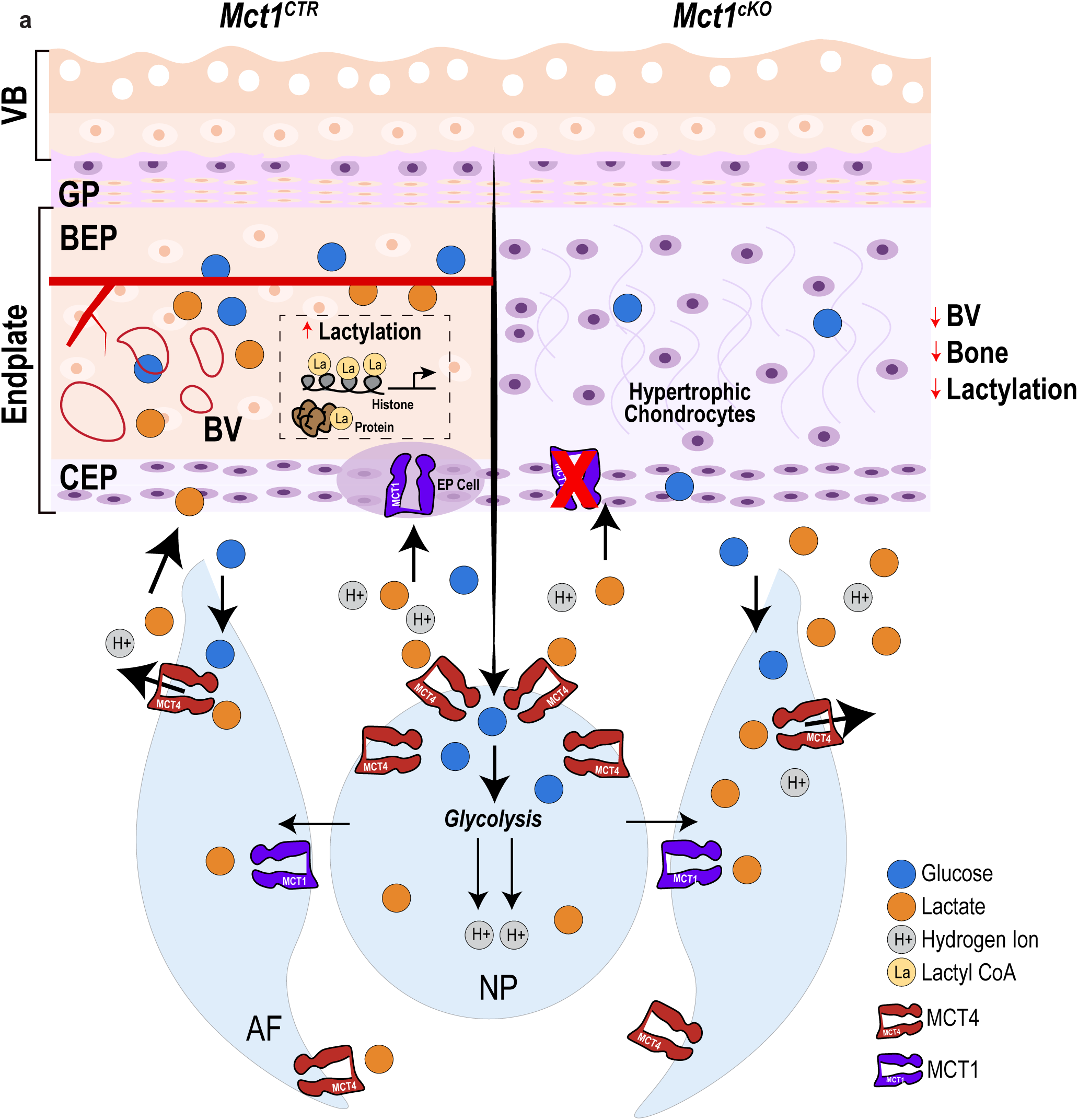
MCT4-MCT1 axis mediates lactate coupling between NP and EP compartments in a growth-dependent manner. a) Schematic representation of MCT-driven lactate coupling between the nucleus pulposus (NP), annulus fibrosus (AF), and endplate (EP), which lies underneath the growth plate (GP) and consists of the bony endplate (BEP) containing blood vessel (BV) and cartilaginous endplate (CEP), in skeletally mature *Mct1^CTR^* and *Mct1^cKO^* mice.

## Discussion

We have uncovered novel insights into the metabolic circuits regulating the function of the intervertebral disc using a conditional mouse model of MCT1 deletion in *Col2a1*-expressing cells (*Mct1^cKO^*). These studies were based on the growing recognition that lactate is more than just a metabolic waste product (21). Instead, lactate has been shown to be a versatile metabolic fuel across various tissue types (16,17). Additionally, lactate functions as a signaling molecule that drives gene expression via histone lactylation (18). Our studies highlight the observation that metabolic cross-talk mediated by lactate-coupling between the disc compartments is essential for the growth and overall health of the spinal motion segment. Our findings reveal that discs of young adult *Mct1^cKO^* mice with MCT1 loss in the EP and AF, tissues derived from the embryonic sclerotome, exhibit clear degenerative changes. We also noted prominent NP degeneration, characterized by fibrosis, cell death, and reduced expression of metabolic health markers involved in glucose homeostasis. However, the absence of *Col2a1*Cre targeting of the NP suggests that these changes were secondary to disrupting lactate homeostasis in the AF and EP. Interestingly, while the outer AF did not appear to be significantly affected by lack of lactate import, the inner AF exhibited lamellar buckling, likely influenced by the substantial structural changes in the NP. As expected, AF showed a higher proportion of thin collagen fibers, indicating increased collagen turnover in the mutants.

The most striking phenotype noted in *Mct1^cKO^* mice was the delayed maturation of endplates and persistence of hypertrophic cartilage. By skeletal maturity, the endplate region develops into two distinct regions: the bony endplate, which overlays a thin, two-cell-thick layer of cartilage that persists throughout life. The bony endplate forms as mature chondrocytes transdifferentiate into bone (5,6). Conversely, in *Mct1^cKO^* mice, the persistence of endplate cartilage was evident by the absence of vascularization, lack of osteogenic markers like TNAP, and reduced mineralization. Furthermore, the robust COL X staining in endplates showed that the persisting cells were hypertrophic chondrocytes. Notably, *Mct1^cKO^* phenocopied *Mct4*-KO mice, wherein NP cells failed to efflux lactate, leading to diminished lactate availability for the endplate chondrocytes (9). Accordingly, these disruptions in lactate-coupling in MCT-mutants resulted in metabolic dysregulation and retardation of endplate differentiation, with chondrocytes unable to progress through the transdifferentiation process into the bone.

We employed spatial transcriptomics to gain deeper insights into the *Mct1^cKO^* phenotype. Notably, the osteoblast cluster in the bony endplate was repositioned to an enthesis-like region in outer AF and at the NP/AF junction. This observation suggests that these cells may acquire an osteoblast-like fate, similar to an observation made in *ank*/*ank* mice (27). The ‘cNP’ and ‘Chondrocyte I’ clusters also showed pronounced changes in their signatures. Interestingly, recent scRNASeq studies showed that the NP cells in mice segregate into two distinct populations, ‘cNP’ and ‘pNP’, a finding consistent with our spatial results (28,29). These clusters showed greater divergence compared to other cell populations within the disc, emphasizing their unique transcriptomic profiles and underscoring their notochordal origin. Analysis of the ‘cNP’ cluster revealed disruptions in circadian clock concepts, which are essential for NP cell adaptation to their specialized niche (30). *Per1* and *Dbp* are ‘clock genes’ directly linked to BMAL1, whose absence is linked to disruptions in ECM homeostasis and pathological endochondral ossification in the disc (31). Changes associated with NP degeneration were also reflected in altered expression of matrix-related genes such as *Acan* and *Sdc4* (32). Additionally, modifications in endocytosis and Golgi apparatus functions, involving genes like *Cavin1* and *Golgb1*, have been implicated in disc degeneration (33,34). Intriguingly, GSEA analysis suggested a shift in the cNP transcriptomic profile toward a muscle-like phenotype, in *Mct1^cKO^* mice characterized by processes such as ‘actin-myosin filament sliding,’ ‘muscle filament sliding,’ and ‘muscle contraction.’ Moreover, the ‘Chondrocyte I” cluster showed strong connectivity between themes relating to *Sarcomeres/Myofibrils Related*, and *BMP Signaling* with genes such as *App, Sost, Col1a1, Col1a2, Fgfr3,* and *Ttn*, indicating changes in muscle-related pathways and consequently, skeletal processes. GSEA further corroborated these outputs, revealing an upregulation in gene expression and metabolic pathways, indicating disruptions in cartilage matrix production and signaling. Recent studies have shown that PPARγ in osteoblasts regulates bone formation and fat mass by regulating sclerostin (SOST) expression; thus, we hypothesize that alterations in sclerostin activity could reflect changes in hypertrophic differentiation (35). Similarly, FGFR3 plays a crucial role in inhibiting chondrocyte proliferation and differentiation, making it a key regulator of bone development. Mutations in *Fgfr3*, as seen in achondroplasia, lead to receptor over-activation, disrupting normal skeletal development and causing disproportionate short stature (36). Furthermore, a study on the *Fgfr3^Y367C^* activating mutation in skeletal stem/progenitor cells showed impaired bone healing, with these cells failing to support cartilage-to-bone transformation, leading to a pseudoarthrosis phenotype (37). These findings suggest that changes within the ‘Chondrocyte I’ cluster could indicate phenotypic alterations or shifts in chondrocyte differentiation. Moreover, the *App* gene that is highly expressed in *Mct1^cKO^* was recently recognized to function as a clusterin protein and may serve as a biomarker for osteoarthritis (38).

When EP chondrocytes were evaluated for bioenergetic profiles, they revealed a predominant reliance on glycolysis. This glycolytic nature remained consistent even after 1-week exposure to lactate in the presence of other metabolic substrates like glutamine, fatty acids, or an osteogenic differentiation protocol. Recently studies by Thompson’s group demonstrated that lactate can suppress cancer cell glycolysis while inducing a metabolic shift towards OXPHOS (16). The *Mct1^cKO^* phenotype raised the possibility that lactate served as a metabolic substrate to drive differentiation. However, given the absence of a strong OXPHOS shift, the mechanism appeared to be distinct from cancer cells or growth plate chondrocytes. ^13^C stable isotope labeling experiments revealed that lactate metabolism contributed to TCA-derived metabolites suggesting that endplate cells rely both on glucose and lactate to replenish the pyruvate pool and utilize both substrates flexibly for TCA intermediate anaplerosis rather than ATP generation.

Previous *in vitro* work by Wang et al. showed that AF cells import lactate with about 10-15% of TCA cycle intermediates being derived from lactate (10). Their conclusion that AF cells metabolize most of the NP-derived lactate is paradoxical in that the AF cells, like the NP cells, reside in an avascular, hypoxic environment and would be expected to generate lactate, and thus, their reliance on this anion as an oxidative substrate is unexpected. To examine this possibility, we performed a [U-^13^C]-glucose and [3-^13^C]-lactate co-labeling experiment and found that glucose predominantly contributed to the pyruvate and citrate pools, indicating that glucose served as the primary substrate, with lactate playing a minor role. Seahorse Flux Analyses further supported the hypothesis that in the presence of lactate, AF cells do not switch to an oxidative phenotype but continue to be glycolytic.

Based on lactate-dependent increases in pan-Kla and H3K18la, without change in pan-Kac, indicated that exogenous lactate could induce lactylation in EP chondrocytes independently of acetylation, which is particularly noteworthy given the broad impact of acetylation on protein function and gene transcription. Recent studies have linked lactylation with histone acetyltransferases (HATs) or lysine acetyltransferases (KATs), such as HBO1 (20) and explored their effect on gene expression (18). Our RNA-Seq analysis revealed a prominent downregulation in fatty acid genes, including *ACSS2*, a recently identified lactyl-CoA synthetase (39), and *KAT5*, the top expressed KAT, along with *KAT2A*, which has been linked to lactyltransferase activity (40). Further informatic analysis revealed themes related to PPARγ, which is required for both adipocyte and cartilage differentiation, in particular its downregulation here potentially playing a critical role in chondrocyte differentiation (41). A recent study of fracture healing showed the importance of PPAR signaling in regulating endochondral bone and vascular development, linking skeletal regeneration to metabolic and transcriptional pathways (42).

Downregulation of *Ankh*, *Enpp2*, and *LPA/Lysophospholipid Signaling* suggested altered pyrophosphate activity. Pyrophosphate is a crucial regulator of bone mineralization as it inhibits hydroxyapatite crystal formation and growth. Moreover, we have shown that the loss of *Ank* causes abnormal mineralization and the transformation of AF cells into an osteoblast-like phenotype (27,43). In terms of lipid metabolism, we observed a predominance of fatty acid-related themes, such as *Fads*, indicating a downregulation in phospholipid or fatty acid metabolism. This observation was further supported by a suppression in cholesterol-related genes *Mgll, Srebf1, Scd,* which are key players in phospholipid metabolism. It should be acknowledged that phospholipids are essential in regulating chondrocyte maturation, notably through the Wnt/β catenin pathway, and in maintaining the integrity of the ECM (44). Beyond structural functions, phospholipids actively participate in signaling processes that drive chondrocyte differentiation and maturation. Of the hormones that regulate cartilage homeostasis and maturation, one of the most important is IGF-1 (45). The observed changes in *Igfbp5*, a classical IGF binding protein, suggests a shift towards altered IGF signaling and subsequently chondrocyte maturation (46). Overall, lactate treatment induces a complex array of changes that contribute to a hypertrophic phenotype, while suppressing the cartilage sclerotomal phenotype.

Computational analysis of the upregulated DEGs indicated changes in growth regulation and development. The upregulation of *Fgfr1* and *Col2a1* suggested a transition from a proliferative to a differentiated state. A previous study demonstrated that FGFR1 signaling in hypertrophic chondrocytes plays a critical role in endochondral bone formation through a complex intracellular mechanism involving neurofibromin (47). Additionally, the upregulation of *Csf1*, *Ccl2*, and *Meox1* may aid in chondrocyte maturation by modulating inflammatory and immune responses that typically drive cell proliferation and migration. This creates a more stable, controlled environment conducive to proper chondrocyte differentiation. In this context, upregulation of excessive growth factor signaling and inflammatory pathways likely aids in the orderly progression of chondrocytes through their maturation stages, contributing to more organized cartilage formation.

In summary, our study reveals a novel role of lactate in growth-dependent metabolic coupling between the endplates and other disc compartments. Beyond its role as a metabolic fuel, lactate acts as a signaling molecule, driving chondrocyte maturation through lactylation. These results highlight the importance of lactate and suggest its potential as a biomarker for normal disc function, warranting further investigation into its role in disc health and disease.

## Materials and Methods

### Mice

All animal studies were approved by the Institutional Animal Care and Use Committee (IACUC) of Thomas Jefferson University. *Mct1* conditional knockout (*Mct1^cKO^*: *Col2a1^CreERT2^Mct1^f/f^*) and control (*Mct1^CTR^*: *Mct1^f/f^*) mice were generated by crossing *Mct1*^f/f^ mice with *Col2a1*^CreERT2^ mice (22,23) (Fig. 1A). For all experiments, 2-week-old female and male mice of all genotypes received an intraperitoneal injection of 100μg/g body weight of tamoxifen (Sigma-Aldrich, St. Louis, MO, USA) dissolved in corn oil (Sigma-Aldrich) for 5 consecutive days to activate Cre recombinase and were analyzed at 6- and 12-months to investigate the effects of *Mct1* loss on disc health. To visualize Cre targeting, a loxP-stop-loxP tdTomato reporter mouse Ai9 (Strain #007909, Jackson Labs) was crossed with mice with (*Col2a1^CreERT2^Mct1^f/f^*) or without (*Col2a1^CreERT2^*) conditional *Mct1* loss.

### Histological Analysis

Dissected spines were immediately fixed in 4% paraformaldehyde for 4 or 24 hours to process for frozen or paraffin embedding, respectively, following decalcification in 20% EDTA at 4°C. 7-μm-thick mid-coronal sections were stained with 1% Safranin-O, 0.05% Fast Green, and 1% Hematoxylin to assess morphology and imaged on an Axio Imager A2 microscope using 5x/0.15 N-Achroplan or 20x/0.5 EC Plan-Neofluar (Carl Zeiss) objectives, Axiocam 105 color camera, and Zen2™ software (Carl Zeiss AG, Germany) (25). Disc degeneration was evaluated using three blinded observers and histological scoring was performed using a Modified Thompson grading scale for the NP and AF, and the Tessier endplate grading scale for the EP (48) histological scores. Histological grades (grades 1-5) increase with severity of degeneration.

Quantification of the percentage of Safranin-O positively stained area in the 12-month endplates (EP) was conducted using the Fuji package of ImageJ. The colors of the images were separated using the “Split channels” function, and manual thresholding was done on the red channel of the images. Compartments of the superior EP and inferior EP were then manually selected using the “Polygon Selection” tool, and the positively stained area was analyzed using the Measure function and Area Fraction measurement.

### TUNEL Staining

The TUNEL assay was performed on disc tissue sections from 12-month-old *Mct1* mice using an “In Situ Cell Death Detection” kit (Roche Diagnostics). Briefly, sections were deparaffinized and permeabilized with Proteinase K (20 μg/mL) for 15 minutes at room temperature. The assay was then conducted following the manufacturer’s instructions, after which the sections were mounted with ProLong® Gold Antifade Mountant containing DAPI (Thermo Fisher Scientific, P36934). Mounted slides were imaged using an Axio Imager 2 microscope, as previously described (25). TUNEL-positive cells and DAPI-positive cells were quantified to assess cell death and cell number in the disc compartments, respectively.

### Picrosirius Red Analysis

Picrosirius Red staining on disc sections of 12-month-old mice was performed to visualize collagen fibril thickness. A polarized light microscope was used to visualize collagen organization, as previously described (25). Under polarized light, collagen fibrils are visualized as green, yellow, or red pixels corresponding to thin, intermediate, or thick fibrils. The color threshold levels were kept constant throughout the analysis.

### Immunohistochemistry

Mid-coronal 7-μm paraffin from 12-month-old mice were deparaffinized in Histoclear and rehydrated in a graded ethanol series before antigen retrieval using one of the following methods: a 30-minute incubation in hot citrate buffer, a 10-minute incubation at room temperature with proteinase K, or a 30-minute incubation at 37°C with Chondroitinase ABC. Following antigen retrieval, the sections were blocked with 10% normal goat serum (Thermo Fisher Scientific, 10,000C) in PBS-T (0.4% Triton X-100 in PBS), and incubated with primary antibodies against MCT1 (1:150; Invitrogen; PA5-72957), GLUT1 (1:100; Abcam; ab115730), MCT4 (1:100; Invitrogen; PA5-87977), COL X (1:500; Abcam; ab58632), EMCN (1:100; Santa Cruz; sc65495), TNAP (1:50; sc271431), COL 1 (1:100; Millipore Sigma; ABT123), ACAN (1:50; Millipore Sigma; AB1031), C.S. (1:300; Abcam; ab11570). Similarly, some discs embedded in OCT were sectioned on CryoStar™ NX70 Cryostat and briefly fixed with acetone. Tissue sections were then blocked with 10% serum (Thermo Fisher Scientific, 10,000C) in PBS-T or M.O.M.™Immunodetection Kit (Vector Laboratories, BMK-2202), and then incubated with primary antibodies against LDHA (1:100; Novus; NBP2-19320), LDHB (1:50; Santa Cruz; sc100775), pan-Kla (1:100, PTM Bio, PTM-1401-RM), and H3K18la (1:100, PTM Bio, PTM-1406RM). Sections were washed and incubated with Alexa Fluor-488 or -594 conjugated secondary antibodies (1:700, Jackson ImmunoResearch, West Grove, PA). for 1 hour at room temperature. In addition, F-CHP (3Helix) assay was performed following the manufacturer’s protocol. Following incubation, sections were washed, mounted, and imaged on Axio Imager 2 microscope (Carl Zeiss) using 5x/0.15 N-Achroplan or 10x/0.3 EC Plan-Neofluar (Carl Zeiss) objectives, X-Cite 120Q Excitation Light Source (Excelitas) and AxioCam MRm R3 camera (Carl Zeiss Microscopy) or Zeiss LSM 800 Axio Inverted confocal microscope with Plan-Apochromat 20x/0.8 and Zen 2 (blue edition) software (Carl Zeiss Microscopy).

### Micro-computed tomography (μCT) Analysis

Spines of 12-month-old *Mct1^CTR^* and *Mct1^cKO^*mice were placed in PBS and scanned on μCT scanner (Skyscan 1272, Bruker, Belgium) at an energy of 50kVp, current of 200μA, and a 10 μm^3^ voxel size resolution. In DataViewer, length of vertebral bones and height of adjacent discs were measured in the dorsal, midline, and ventral regions of the sagittal plane and averaged; from this, disc height index (DHI) was calculated as previously described (25). Cortical and trabecular bone microstructure was also analyzed from these scans and the following parameters were analyzed: trabecular separation (Tb. Sp.), trabecular thickness (Tb. Th.), trabecular number (Tb. N.), trabecular bone volume fraction (BV/TV), and trabecular bone mineral density (Trab. BMD). Cortical bone mineral density (Cort. BMD), mean cross-sectional bone thickness (Cs. Th.), cortical BV, and mean cross sectional bone area for the cortical bone. The bone mineral density (BMD) of the caudal endplates was tabulated in a region of interest (ROI) defined by outlining the superior and inferior bony endplates that lie below the growth plate, as previously described (27).

### Spatial Transcriptomics

FFPE blocks of *Mct1^CTR^* and *Mct1^cKO^*mice were generated, embedding one caudal (Ca7/8) level from four different mice into each paraffin block. Following RNA quality assessment (DV200 > 30), 5-μm sections were cut and placed onto Nexterion Slide H – 3D Hydrogel Coated slides. Hematoxylin and Eosin (H&E) staining was performed to orient the tissue sections. The slides were loaded into the Visium CytAssist system to facilitate the transfer of oligonucleotide barcodes from the Visium slides to the tissue sections. The Visium CytAssist Gene Expression for FFPE workflow was employed to automate the transfer of transcriptomic probes from glass slides to Visium slides. Tissue permeabilization was optimized by increasing the decrosslinking temperature from 70°C to 95°C to enhance probe access to genomic DNA (gDNA). This adjustment improved the detection of Unique Molecular Identifier (UMI) counts from both gDNA and mRNA probe ligation events. Following barcode transfer, the barcodes were collected, and sequencing libraries were prepared using the NovaSeq-Sp100 kit, following the manufacturer’s protocol.

Raw reads from Visium libraries were processed using SpaceRanger v3.0.0. Associated CytAssist and H&E images were manually aligned using Loupe v8.0.0. Raw gene counts from 8 μm bins were further processed using Seurat v5.0.3. Bins with less than 20 counts or 20 observed genes were discarded. To form bins larger than 16 μm, an array with the coordinates and gene expression values from the 16 μm binned data was convolved using a matrix of ones and size equal to the desired binning multiple. Then, rows and columns were dropped such that none of the convolved 16 μm bins overlap. Associated spatial metadata was recalculated by linear transformations of the 16 μm metadata using the combining multiplier. For this analysis, 48 μm bins were generated and clusters were identified using the FindClusters function in Seurat with default parameters and resolution=1.0. Cluster annotation was manually performed by analyzing the top marker DEGs in Seurat for each cluster.

Cluster-specific differential gene expression analysis was conducted across control and experimental conditions with spatial spots aggregated over all replicates. Genes were filtered for having at least 10% of spatial spots expressed in one condition. Statistically significant differentially expressed genes (DEGs) were found using the FindMarkers function in Seurat v5.0.3 with MAST, with q<0.05 after FDR correction for multiple comparisons. Pathway analyses were performed for DEGs per cluster using the prerank function on average log2 fold change with default parameters in GSEApy v1.1.3. The GO Biological Processes 2021 gene set library was used as reference. Significant overlap of ranked DEGs with pathways was depicted both at q-value of 0.25 as recommended by GSEA documentation and at q-value of 0.05, after FDR correction. The normalized enrichment score was used to assess directionality of association between DEGs and pathways - a positive score denotes the genes at the top of the ranked list (upregulated genes) as having significant overlap with the pathway, whereas a negative score corresponds to the bottom of the ranked list (downregulated genes). Spatial sequencing data is deposited in the GEO database (GSE287499).

### CompBio Analysis

Spatial transcriptomics data from *Mct1^CTR^* and *Mct1^cKO^* mice, along with bulk RNA-Seq data were analyzed using the GTAC-CompBio Analysis Tool (PercayAI Inc., St. Louis, MO). CompBio employs the Biological Knowledge Generation Engine (BKGE) to extract PubMed abstracts referencing input differentially expressed genes (DEGs) and identify relevant biological processes and pathways. Conditional probability analysis was applied to compute enrichment scores, which were normalized against randomized query groups to determine statistical significance. Related concepts derived from DEGs are grouped into broader, higher-level themes, which represent overarching biological patterns, such as pathways, processes, cell types, and structures. The Normalized Enrichment Score (NEScore) quantifies enrichment magnitude, and results were clustered based on fold enrichment and rarity as previously described (25).

### Primary AF and Endplate Cell Isolation

Primary AF cells were harvested and cultured from adult Sprague Dawley rats. AF cells were supplemented with DMEM containing 10% FBS. To investigate the effects of lactate on cell metabolism (49), AF cells were cultured at 5% O_2_ in a hypoxia workstation (InvivO_2_ 300; Baker Ruskinn, USA) with or without sodium L-lactate for 24 hours. Primary rat EP cells were harvested and cultured from 3-week-old Sprague Dawley rats. For the cell differentiation studies, EP cells were cultured in complete DMEM with 10 mM β-glycerophosphate (Sigma-Aldrich, G9422), 50 μg/ml L-ascorbate-2-phosphate, and 1x insulin, transferrin, selenium (Gibco, 41400045) and varying concentrations sodium L-lactate up to 7 days under 5% O_2_ (InvivO_2_ 300; Baker Ruskinn, USA). To measure protein lactylation, EP cells were cultured in DMEM supplemented with 20 mM sodium lactate or 20 mM NaCl for 24 hours.

### Glucose and Lactate Stable Isotope Tracing Analysis

Primary AF (n = 4 sets/condition) and EP (n = 5 sets/condition) cells were plated in 6 cm dishes at a density of ∼10^6^ cells and incubated in 5% O_2_ for 24 hours prior to metabolomic analysis the following day. All experiments were completed in DMEM media without glucose, glutamine, phenol red, sodium pyruvate, and sodium bicarbonate (Sigma, D5030) that was supplemented with 44 mM sodium bicarbonate, 4 mM glutamine, 5 mM glucose, and 10% dialyzed FBS (Corning 35-071-CV). The next day, the cultures were replenished with media containing 5 mM [U-^13^C] glucose (Cambridge Isotope Laboratories, CLM-1396) and varying concentrations up to 10 mM [3-^13^C] lactate (Cambridge Isotope, CLM-1579) as isotope tracers and incubated for 30 mins to limit the lactate contribution to the second turn of the TCA cycle. Another experiment (n = 5 sets) was conducted in EP cells treated with 10 mM [3-^13^C] lactate and 5 mM unlabeled glucose for 24 hours in hypoxia to determine where the lactate pool accumulates. Cells were harvested and polar metabolites were extracted by the addition of 1 ml ice-cold extraction solution of 80% methanol/20% water. Unlabeled samples were also prepared for each experiment, from which an unlabeled sample pool was generated. Metabolomics analyses were performed by The Wistar Institute Proteomics and Metabolomics Shared Resource. Samples were analyzed by LC-MS/MS on Thermo Scientific Q Exactive HF-X or Plus mass spectrometers interfaced with Vanquish UHPLC Systems. Samples were analyzed in a pseudorandomized order.

LC separation was performed using a ZIC-pHILIC column, 150 × 2.1 mm, 5 μM (EMD Millipore) maintained at 45 °C. Mobile phase A was 20 mM ammonium carbonate, 5 µM medronic acid, 0.1% ammonium hydroxide, pH 9.2, and mobile phase B was acetonitrile. Analytical separation was performed at 0.2 ml/min flow rate using the following step-wise gradient: 0 min, 85% B; 2 min, 85% B; 17 min, 20% B; 17.1 min, 85% B; and 26 min, 85% B. Samples were analyzed by either Full MS scans with polarity switching (all samples) or Full MS scans with data-dependent MS/MS scans with separate acquisitions for positive and negative polarities (unlabeled sample pool). Relevant MS parameters include: sheath gas, 40; auxiliary gas, 10; sweep gas, 2; auxiliary gas heater temperature, 350°C; spray voltage, 3.5/3.2 kV for positive/negative polarities; capillary temperature, 325°C; S-lens RF, 40 for HF-X and 50 for Plus. Full MS scans were acquired using a scan range of 65 to 975 m/z; resolution of 120,000 for HF-X and 70,000 for Plus; automated gain control (AGC) target of 1E6; and maximum injection time (IT) of 100 ms. Data-dependent MS/MS was performed on top 10 ions; resolution of 15,000 for HF-X and 17,500 for Plus; AGC target of 5E4, maximum IT of 50 ms, isolation width of 1.0 m/z, and a stepped normalized collision energy of 20, 40, 60.

Raw data were analyzed using Compound Discoverer 3.3 SP3 (Thermo Scientific). Metabolites were identified in the unlabeled sample pool by accurate mass in conjunction with either retention time based on pure standards or MS/MS fragmentation by querying the mzCloud database. These identifications were applied to labeled samples with consideration for all possible ^13^C isotopologues. Metabolite levels were quantified by integrating peak areas from Full MS data, and data were corrected for natural ^13^C abundance and tracer purity (50).

### Seahorse Metabolic Assays

Primary rat AF or EP cells were plated in Seahorse microplates at appropriate densities (15,000 cells/well for 24-hour experiments and 4,000 cells/well for long-term experiments) and were allowed to adhere overnight to 2 days. Cell culture media was then replenished with or without sodium L-lactate under 5% O_2_ for 24 hours, or up to 7 days prior to Seahorse measurements. Prior to analysis, cells were washed twice with Krebs-Ringer-Phosphate-Hepes (KRPH) buffer and incubated in KRPH + 0.1% BSA with or without lactate. The microplates were degassed for 1 hour before placing the cells in a Seahorse Flux Analyzer. For ATP consumption assays, cells are sequentially injected with 5 mM glucose, 2 μg/ml oligomycin, 1 μΜ rotenone with 1μΜ myxothiazol while monitoring changesin oxygen consumption rate (OCR) and extracellular acidification rate (ECAR). For substrate utilization assays, cells were sequentially injected with 5 mM glucose with 4 mM glutamine or pamitate-BSA (palmitate concentration 150 μM); 2 μg/ml oligomycin, 1 μΜ FCCP followed by 5 μΜ BPTES (GLS inhibitor) or 5 μΜ Etomoxir (CPT1 inhibitor). Raw ECAR and OCR raw traces were plotted and used to quantify ATP production – specifically, the partitioning of glycolytic and oxidative ATP (51,52). Parameters relating to glycolysis (glucose stimulated minus basal), glycolytic capacity (after oligomycin addition), and reserved capacity (oligomycin-stimulated minus basal) were extracted from ECAR values. OCR values were used to determine ATP-coupled respiration (after oligomycin addition), endogenous (no substrate), and basal (after substrate addition) as previously described (1). Results were normalized to total protein.

### Western Blot

Treated and untreated EP cells were lysed, and 35 μg of whole cell lysate or 15 μg of purified histones (Abcam, ab113476), were electroblotted to PVDF membranes (EMD Millipore, IPVH00010). The membranes were blocked with 4% (w/v) nonfat dry milk in TBS-T and incubated overnight at 4°C with antibodies against pan-Kla (1:1000; PTM-1401) pan-Kac (1:1000; PTM-101), H3K18la (1:1000; PTM-1406RM), H3K27ac (1:1000; PTM-116RM), and housekeeping antibodies against total Histone H3 (1:3000; CS; 4499). An ECL reagent (Prometheus, 20-301) was utilized to detect immunolabeling using an Azure 300 system. Densitometric analysis was conducted using ImageJ software.

### RNA-seq

DNA-free RNA was isolated from treated endplate cells using a RNeasy^®^ Micro kit (Qiagen, Venlo, Netherlands). RNA quality and concentration were assessed using a Nanodrop ND 100 spectrophotometer (Thermo Fisher Scientific) and an Agilent 2200 Tape Station (Agilent Technologies, Palo Alto, CA, USA) respectively. cDNA was synthesized using the RNA to cDNA EcoDry Premix (Takara, #639549) and subsequently used for qPCR validation. For bulk RNA sequencing, purified RNA was sent to Azenta Life Sciences/Genewiz (Chelmsford, MA, USA). Library preparation, including poly(A) enrichment and cDNA synthesis, was performed by Azenta according to their standard protocols. Sequencing was conducted on an Illumina platform, generating paired-end reads with a length of 150 bp. Raw sequencing data underwent quality control and were provided as FASTQ files for bioinformatic analysis. Differential gene expression was performed using a threshold of an adjusted p-value (FDR) < 0.05 and log2 fold change cutoff to identify significant changes in expression. Spatial sequencing data is available in the GEO database (GSE287056).

### Immunocytochemistry

EP cells were plated on collagen I-coated glass coverslips and treated for 24 hours under hypoxic conditions. Cells were then fixed with ice cold methanol for 10 min., permeabilized with 0.1% Triton X-100 for 15 min. and blocked with 1% BSA for 1 hour. Cells were incubated overnight at 4°C with primary antibodies against H3K18la (1:1000; PTM-1406RM) and H3K27ac (1:1000; PTM-116RM), diluted in blocking buffer. After washing, cells were incubated with Alexa Fluor-594-conjugated secondary antibodies and mounted with ProLong Gold Antifade Mountant with DAPI. Negative controls were used to confirm the specificity of staining. Cells were visualized using a Zeiss LSM 800 Axio Inverted confocal microscope (Plan-Apochromat 40x/1.3 oil or 63x/1.40 oil).

### Statistical Analysis

Statistical analysis was performed using Prism 10 (GraphPad, La Jolla, CA) with data presented as mean +/- standard error mean to represent the precision of the mean estimate. Normality was assessed using the Shapiro-Wilk tests to evaluate differences between distributions. For normally distributed data, an unpaired t-test was used for comparisons, while the Mann-Whitney U test was applied for non-normally distributed data. To compare multiple distributions of non-normally distributed data, the Kruskal-Wallis test with Dunn’s multiple comparison was used. For normally distributed data, one-way ANOVA with Sidak’s post-hoc multiple comparison test was used. The distributions of Modified Thompson Grading and fiber thickness data were analyzed using a Fisher’s exact test or χ² test with a significance level set at 0.05. All tests were two-tailed tests. F- and t- statistics and degrees of freedom for ANOVA and t-tests were included in the Fig. Legends, accordingly.

## Data Availability

Spatial transcriptomics and bulk RNA-Seq data associated with this study are deposited in the GEO database under the accession codes: GSE287499 and GSE287056, respectively. All generated and analyzed data are included in this publication.

## Acknowledgments

This work was supported by grants from the NIH/NIAMS R01AR064733, R01AR074813, and 2R56 AR055655-16A1 to MVR. We thank Mallory Toci, Bharvi Chavre, Cindy Pham and Mei Smyers for technical help with experiments. We also thank Drs. Regis O’Keefe and Jeffrey Rothstein, Brett Morrison for providing the mouse models used in this study. We acknowledge 10x Genomics in providing consultations that facilitated aspects of this work. Funding support for The Wistar Institute Proteomics and Metabolomics Shared Resource was provided by Cancer Center Support Grant P30 CA010815. The Thermo Q Exactive HF-X mass spectrometer used in this study was purchased with the National Institutes of Health instrument award S10 OD023586.

## Author Contributions

MT and MVR have designed experiments. MT and WZ conducted experiments. MT, ARG, and KT performed data analysis. MT and MVR wrote and edited the manuscript, which was approved by all authors.

---

<u>Supplementary Figure 1</u>. **MCT1 loss affects endplate mineralization but does not markedly impact vertebral bone.** a) μCT scan of a caudal vertebral motion segment of 12-month-old *Mct1^CTR^* and *Mct1^cKO^* mice, scale bar = 1 mm. b) Disc height, vertebral length, and disc height index (DHI). *n* = 10 mice/genotype (CTR: 6 f, 4 m; cKO: 5 f, 5 m), Ca7/8, 8/9/animal were assessed. c) Trabecular and cortical bone morphology analysis showing bone volume fraction (BV/TV), trabecular thickness (Trab. Th.), separation (Trab. Sp.), number (Trab. N.), and bone mineral density (Trab. BMD), cortical bone volume (BV), cross-sectional thickness, mean cross-sectional bone area and cortical BMD. *n* = 4 animals/genotype (2 m, 2 f); Ca7-Ca8 /animal were assessed. d) Representative μCT images of superior and inferior endplates of Ca7/8 disc and endplate BMD analysis. *n* = 10 mice/genotype (CTR: 6 f, 4 m; cKO: 5 f, 5 m), 2 levels; 4 endplates/mouse were assessed. Scale bar = 125 μm. Data represented as violin or box and whisker plots with median and quartiles. Mann Whitney test or t-test as appropriate was used to test significance.

<u>Supplementary Figure 2</u>. **Spatial transcriptomics shows altered gene expression in different disc cell clusters of *Mct1^cKO^* mice.** Feature plots showing spatial expression for specific marker genes in (a) NP (b) AF (c) chondrocyte, and (d) osteoblast clusters in *Mct1^cKO^* and *Mct1^CTR^*spinal motion segments, *n* = 4 mice/genotype; 2 m, 2 f.

<u>Supplementary Figure 3</u>. ***Mct1^cKO^* mice show altered transcriptomic profiles.** a) Volcano plots showing downregulated (green) and upregulated (pink) DEGs in the ‘central NP’ cluster of *Mct1^cKO^*. b) Gene set enrichment analysis (GSEA) of DEGs in ‘cNP’ cluster from *Mct1^cKO^* mice. The blue dotted line represents a q-value of 0.25, while the black dotted line indicates a more stringent q-value of 0.05. c) Violin plots show the distributions of DEGs within the ‘cNP’ cluster. d) Volcano plots of downregulated-(green) and upregulated (pink) DEGs in the ‘Chondrocyte I’ cluster from *Mct1^cKO^* mice. e) GSEA analysis of DEGs in the ‘Chondrocyte I’ cluster of *Mct1^cKO^*. f) Violin plots showing the distribution of DEGs in the ‘Chondrocyte I’ cluster of cKO mice (n = 4 mice/genotype; 2 m, 2 f).

<u>Supplementary Figure 4</u>. **MCT1 deletion disrupts the chondrocyte gene expression.** a) Thematic organization of concepts generated by CompBio, derived from a literature analysis of DEGs in the ‘Chondrocyte I’ cluster in the *Mct1^cKO^* mouse discs, highlighting key physiological features impacted by disrupted lactate uptake. Key superclusters of themes include muscle-related pathways, skeletal maturation, actin-cytoskeleton organization and AKT/PI3K/MAPK signaling (green) and rRNA processing and mitochondrial lrRNA export (red). b) DEGs associated with key themes plotted according to their CompBio entity high score (4 mice/genotype; 2 m, 2 f).

<u>Supplementary Figure 5</u>. **AF cells are primarily glycolytic**. Effect of 24-hour lactate supplementation on bioenergetics in AF cells. a) Schematic representation of dual [U-^13^C] glucose and [3-^13^C] lactate tracing, illustrating carbon flow into the TCA cycle. b) Percent labeling of citrate and pyruvate after 24 hr from 5 mM [U-^13^C] glucose with increasing concentrations of labeled lactate (up to 10 mM) in AF cells. Data are presented as the mean ± SEM, *n* = 4 independent samples/condition) Normalized traces of ECAR and OCR measurements, with corresponding (d) quantification of OCR parameters including endogenous, basal, ATP-coupled OCR, and ECAR parameters, including glycolysis, glycolytic capacity and reserved capacity normalized to the no-lactate group. Data is shown as violin plots with median and interquartile ranges. (e) The effect of lactate supplementation on ATP production rate in AF cells (*n* = 5 experiments, with 6 technical repeats/group/trace). Significance was determined using One-way ANOVA with Sidak’s or Dunnett’s post hoc test.

<u>Supplementary Figure 6</u>. **Lactate supplementation induces transcriptional reprogramming in endplate cells.** (a) Hierarchical clustering of differentially expressed genes (FDR p < 0.05) in lactate-treated endplate cells. (b) Volcano plots of down- (green) and up- (pink) regulated DEGs in endplate cells with lactate treatment (c) Hierarchical clustering of curated subset of lactyl-specific DEGs in lactate-treated vs control endplate cells plotted as transcripts per million (TPM). (*n* = 5 experiments; 2 conditions: 20 mM lactate or NaCl)

<u>Supplementary Figure 7</u>. **Lactate-treated EP cells upregulate transcriptional programs related to growth regulation and development.** (a) CompBio-generated thematic organization of concepts, derived from literature analysis of differentially upregulated genes in lactate-treated EP cells, reveals key physiological features related to chemotaxis (pink), hormone feedback regulation (blue), and skeletal development (teal) clusters. (b) Top DEGs enriched in key themes are displayed as entities based on their CompBio entity high score, (FDR p < 0.05), (*n* = 5 experiments; 2 conditions: 20 mM lactate or NaCl).

## References

1. Johnston SN, Silagi ES, Madhu V, Nguyen DH, Shapiro IM, Risbud MV. GLUT1 is redundant in hypoxic and glycolytic nucleus pulposus cells of the intervertebral disc. JCI Insight. 2023 Apr 24;8(8).

2. Agrawal A, Guttapalli A, Narayan S, Albert TJ, Shapiro IM, Risbud MV. Normoxic stabilization of HIF-1alpha drives glycolytic metabolism and regulates aggrecan gene expression in nucleus pulposus cells of the rat intervertebral disk. Am J Physiol, Cell Physiol. 2007 Aug;293(2):C621–31.

3. Silagi ES, Schipani E, Shapiro IM, Risbud MV. The role of HIF proteins in maintaining the metabolic health of the intervertebral disc. Nat Rev Rheumatol. 2021 Jul;17(7):426– 39.

4. Chen N, Wu RWH, Lam Y, Chan WCW, Chan D. Hypertrophic chondrocytes at the junction of musculoskeletal structures. Bone Rep. 2023 Dec;19:101698.

5. Marcucio RS, Miclau T, Bahney CS. A Shifting Paradigm: Transformation of Cartilage to Bone during Bone Repair. J Dent Res. 2023 Jan;102(1):13–20.

6. Hu DP, Ferro F, Yang F, Taylor AJ, Chang W, Miclau T, et al. Cartilage to bone transformation during fracture healing is coordinated by the invading vasculature and induction of the core pluripotency genes. Development. 2017 Jan 15;144(2):221–34.

7. Halestrap AP. The SLC16 gene family - structure, role and regulation in health and disease. Mol Aspects Med. 2013 Jun;34(2–3):337–49.

8. Dimmer KS, Friedrich B, Lang F, Deitmer JW, Bröer S. The low-affinity monocarboxylate transporter MCT4 is adapted to the export of lactate in highly glycolytic cells. Biochem J. 2000 Aug 15;350 Pt 1:219–27.

9. Silagi ES, Novais EJ, Bisetto S, Telonis AG, Snuggs J, Le Maitre CL, et al. Lactate efflux from intervertebral disc cells is required for maintenance of spine health. J Bone Miner Res. 2020 Mar;35(3):550–70.

10. Wang D, Hartman R, Han C, Zhou C-M, Couch B, Malkamaki M, et al. Lactate oxidative phosphorylation by annulus fibrosus cells: evidence for lactate-dependent metabolic symbiosis in intervertebral discs. Arthritis Res Ther. 2021 May 21;23(1):145.

11. Pascale RM, Calvisi DF, Simile MM, Feo CF, Feo F. The Warburg Effect 97 Years after Its Discovery. Cancers (Basel). 2020 Sep 30;12(10).

12. Halestrap AP, Meredith D. The SLC16 gene family-from monocarboxylate transporters (MCTs) to aromatic amino acid transporters and beyond. Pflugers Arch. 2004 Feb;447(5):619–28.

13. Hollander JM, Li L, Rawal M, Wang SK, Shu Y, Zhang M, et al. A critical bioenergetic switch is regulated by IGF2 during murine cartilage development. Commun Biol. 2022 Nov 11;5(1):1230.

14. Stegen S, Rinaldi G, Loopmans S, Stockmans I, Moermans K, Thienpont B, et al. Glutamine metabolism controls chondrocyte identity and function. Dev Cell. 2020 Jun 8;53(5):530–544.e8.

15. Pérez-Escuredo J, Dadhich RK, Dhup S, Cacace A, Van Hée VF, De Saedeleer CJ, et al. Lactate promotes glutamine uptake and metabolism in oxidative cancer cells. Cell Cycle. 2016;15(1):72–83.

16. Cai X, Ng CP, Jones O, Fung TS, Ryu KW, Li D, et al. Lactate activates the mitochondrial electron transport chain independently of its metabolism. Mol Cell. 2023 Nov 2;83(21):3904–3920.e7.

17. Hui S, Ghergurovich JM, Morscher RJ, Jang C, Teng X, Lu W, et al. Glucose feeds the TCA cycle via circulating lactate. Nature. 2017 Nov 2;551(7678):115–8.

18. Zhang D, Tang Z, Huang H, Zhou G, Cui C, Weng Y, et al. Metabolic regulation of gene expression by histone lactylation. Nature. 2019 Oct 23;574(7779):575–80.

19. Merkuri F, Rothstein M, Simoes-Costa M. Histone lactylation couples cellular metabolism with developmental gene regulatory networks. Nat Commun. 2024 Jan 2;15(1):90.

20. Niu Z, Chen C, Wang S, Lu C, Wu Z, Wang A, et al. HBO1 catalyzes lysine lactylation and mediates histone H3K9la to regulate gene transcription. Nat Commun. 2024 Apr 26;15(1):3561.

21. Brooks GA. Lactate as a fulcrum of metabolism. Redox Biol. 2020 Aug;35:101454.

22. Philips T, Mironova YA, Jouroukhin Y, Chew J, Vidensky S, Farah MH, et al. MCT1 Deletion in Oligodendrocyte Lineage Cells Causes Late-Onset Hypomyelination and Axonal Degeneration. Cell Rep. 2021 Jan 12;34(2):108610.

23. Chen M, Lichtler AC, Sheu T-J, Xie C, Zhang X, O’Keefe RJ, et al. Generation of a transgenic mouse model with chondrocyte-specific and tamoxifen-inducible expression of Cre recombinase. Genesis. 2007 Jan;45(1):44–50.

24. Song F, Lee WD, Marmo T, Ji X, Song C, Liao X, et al. Osteoblast-intrinsic defect in glucose metabolism impairs bone formation in type II diabetic mice. BioRxiv. 2023 Jan 18;

25. Tsingas M, Ottone OK, Haseeb A, Barve RA, Shapiro IM, Lefebvre V, et al. Sox9 deletion causes severe intervertebral disc degeneration characterized by apoptosis, matrix remodeling, and compartment-specific transcriptomic changes. Matrix Biol. 2020 Dec;94:110–33.

26. Ottone OK, Mundo JJ, Kwakye BN, Slaweski A, Collins JA, Wu Q, et al. Oral citrate supplementation mitigates age-associated pathological intervertebral disc calcification in LG/J mice. BioRxiv. 2024 Jul 19;

27. Ohnishi T, Tran V, Sao K, Ramteke P, Querido W, Barve RA, et al. Loss of function mutation in Ank causes aberrant mineralization and acquisition of osteoblast-like-phenotype by the cells of the intervertebral disc. Cell Death Dis. 2023 Jul 19;14(7):447.

28. Tan Z, Chen P, Dong X, Guo S, Leung VYL, Cheung JPY, et al. Progenitor-like cells contributing to cellular heterogeneity in the nucleus pulposus are lost in intervertebral disc degeneration. Cell Rep. 2024 Jun 25;43(6):114342.

29. Zhang C, Zhong L, Lau YK, Wu M, Yao L, Schaer TP, et al. Single cell RNA sequencing reveals emergent notochord-derived cell subpopulations in the postnatal nucleus pulposus. FASEB J. 2024 Jan;38(1):e23363.

30. Suyama K, Silagi ES, Choi H, Sakabe K, Mochida J, Shapiro IM, et al. Circadian factors BMAL1 and RORα control HIF-1α transcriptional activity in nucleus pulposus cells: implications in maintenance of intervertebral disc health. Oncotarget. 2016 Apr 26;7(17):23056–71.

31. Dudek M, Morris H, Rogers N, Pathiranage DR, Raj SS, Chan D, et al. The clock transcription factor BMAL1 is a key regulator of extracellular matrix homeostasis and cell fate in the intervertebral disc. Matrix Biol. 2023 Sep;122:1–9.

32. Sao K, Risbud MV. Sdc4 deletion perturbs intervertebral disc matrix homeostasis and promotes early osteopenia in the aging mouse spine. Matrix Biol. 2024 Aug;131:46–61.

33. Risbud M, Madhu V, Hernandez-Meadows M, Coleman A, Sao K, Inguito K, et al. The loss of OPA1 accelerates intervertebral disc degeneration and osteoarthritis in aged mice. Res Sq. 2024 Feb 20;

34. Bach FC, Zhang Y, Miranda-Bedate A, Verdonschot LC, Bergknut N, Creemers LB, et al. Increased caveolin-1 in intervertebral disc degeneration facilitates repair. Arthritis Res Ther. 2015;18(1):59.

35. Kim SP, Seward AH, Garcia-Diaz J, Alekos N, Gould NR, Aja S, et al. Peroxisome proliferator activated receptor-γ in osteoblasts controls bone formation and fat mass by regulating sclerostin expression. iScience. 2023 Jul 21;26(7):106999.

36. Shazeeb MS, Cox MK, Gupta A, Tang W, Singh K, Pryce CT, et al. Skeletal Characterization of the Fgfr3 Mouse Model of Achondroplasia Using Micro-CT and MRI Volumetric Imaging. Sci Rep. 2018 Jan 11;8(1):469.

37. Julien A, Perrin S, Duchamp de Lageneste O, Carvalho C, Bensidhoum M, Legeai-Mallet L, et al. FGFR3 in Periosteal Cells Drives Cartilage-to-Bone Transformation in Bone Repair. Stem Cell Reports. 2020 Oct 13;15(4):955–67.

38. Kovács P, Pushparaj PN, Takács R, Mobasheri A, Matta C. The clusterin connectome: Emerging players in chondrocyte biology and putative exploratory biomarkers of osteoarthritis. Front Immunol. 2023 Mar 15;14:1103097.

39. Zhu R, Ye X, Lu X, Xiao L, Yuan M, Zhao H, et al. ACSS2 acts as a lactyl-CoA synthetase and couples KAT2A to function as a lactyltransferase for histone lactylation and tumor immune evasion. Cell Metab. 2024 Nov 12;

40. Sun W, Jia M, Feng Y, Cheng X. Lactate is a bridge linking glycolysis and autophagy through lactylation. Autophagy. 2023 Dec;19(12):3240–1.

41. Christofides A, Konstantinidou E, Jani C, Boussiotis VA. The role of peroxisome proliferator-activated receptors (PPAR) in immune responses. Metab Clin Exp. 2021 Jan;114:154338.

42. Grimes R, Jepsen KJ, Fitch JL, Einhorn TA, Gerstenfeld LC. The transcriptome of fracture healing defines mechanisms of coordination of skeletal and vascular development during endochondral bone formation. J Bone Miner Res. 2011 Nov;26(11):2597–609.

43. Novais EJ, Narayanan R, Canseco JA, van de Wetering K, Kepler CK, Hilibrand AS, et al. A new perspective on intervertebral disc calcification-from bench to bedside. Bone Res. 2024 Jan 22;12(1):3.

44. Dao DY, Jonason JH, Zhang Y, Hsu W, Chen D, Hilton MJ, et al. Cartilage-specific β-catenin signaling regulates chondrocyte maturation, generation of ossification centers, and perichondrial bone formation during skeletal development. J Bone Miner Res. 2012 Aug;27(8):1680–94.

45. Wang Y, Cheng Z, Elalieh HZ, Nakamura E, Nguyen M-T, Mackem S, et al. IGF-1R signaling in chondrocytes modulates growth plate development by interacting with the PTHrP/Ihh pathway. J Bone Miner Res. 2011 Jul;26(7):1437–46.

46. Bobola N, Engist B. IGFBP5 is a potential regulator of craniofacial skeletogenesis. Genesis. 2008 Jan;46(1):52–9.

47. Karolak MR, Yang X, Elefteriou F. FGFR1 signaling in hypertrophic chondrocytes is attenuated by the Ras-GAP neurofibromin during endochondral bone formation. Hum Mol Genet. 2015 May 1;24(9):2552–64.

48. Tessier S, Tran VA, Ottone OK, Novais EJ, Doolittle A, DiMuzio MJ, et al. TonEBP-deficiency accelerates intervertebral disc degeneration underscored by matrix remodeling, cytoskeletal rearrangements, and changes in proinflammatory gene expression. Matrix Biol. 2020 May;87:94–111.

49. Wang N, Jiang X, Zhang S, Zhu A, Yuan Y, Xu H, et al. Structural basis of human monocarboxylate transporter 1 inhibition by anti-cancer drug candidates. Cell. 2021 Jan 21;184(2):370–383.e13.

50. Di Marcantonio D, Martinez E, Kanefsky JS, Huhn JM, Gabbasov R, Gupta A, et al. ATF3 coordinates serine and nucleotide metabolism to drive cell cycle progression in acute myeloid leukemia. Mol Cell. 2021 Jul 1;81(13):2752–2764.e6.

51. Mookerjee SA, Gerencser AA, Nicholls DG, Brand MD. Quantifying intracellular rates of glycolytic and oxidative ATP production and consumption using extracellular flux measurements. J Biol Chem. 2017 Apr 28;292(17):7189–207.

52. Mookerjee SA, Nicholls DG, Brand MD. Determining maximum glycolytic capacity using extracellular flux measurements. PLoS ONE. 2016 Mar 31;11(3):e0152016.

