## Supplementary Figures for "Lactate metabolic coupling between the endplates and nucleus pulposus via MCT1 is essential for intervertebral disc health"

Supplementary Figure 1

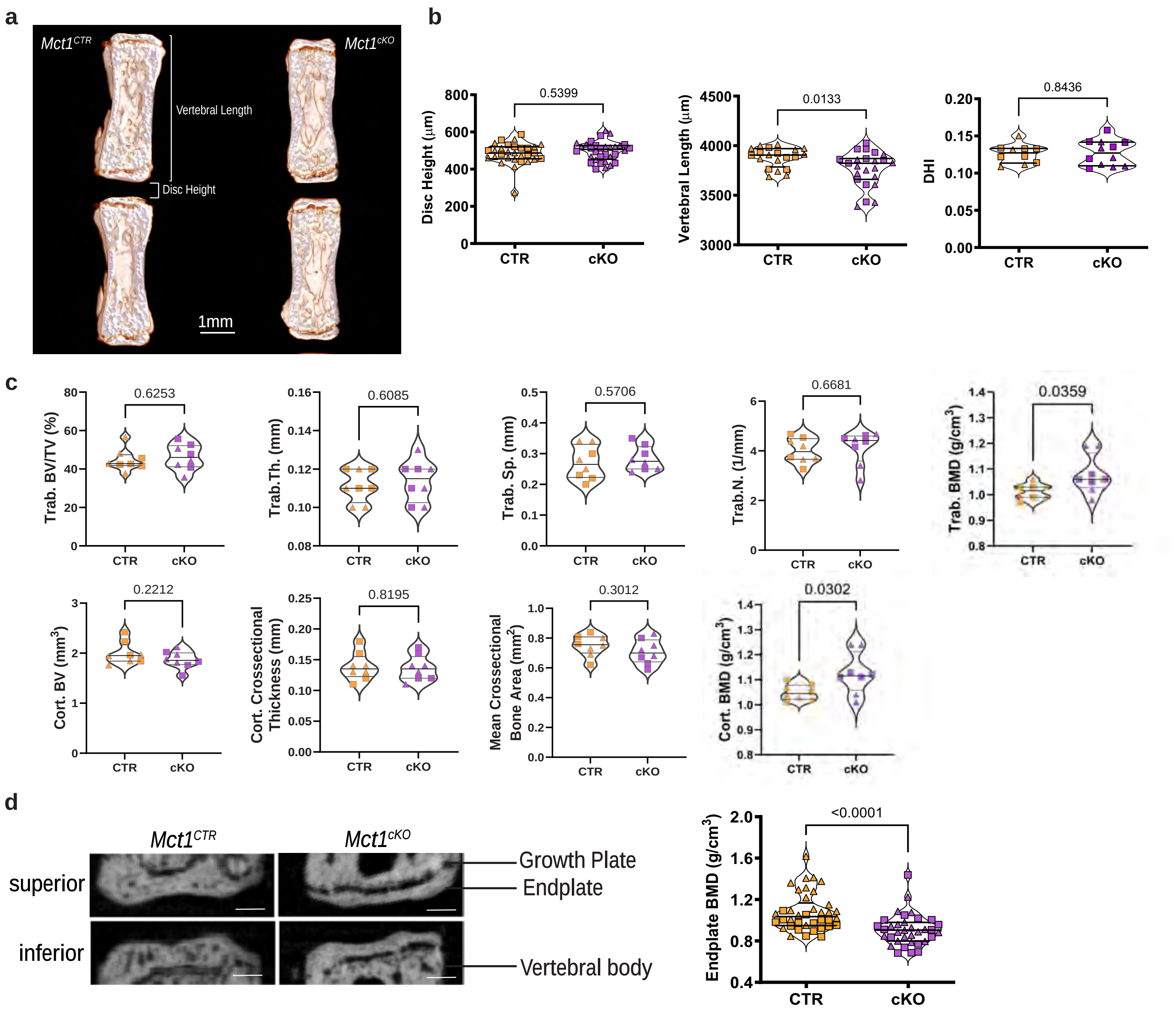

Supplementary Figure 2

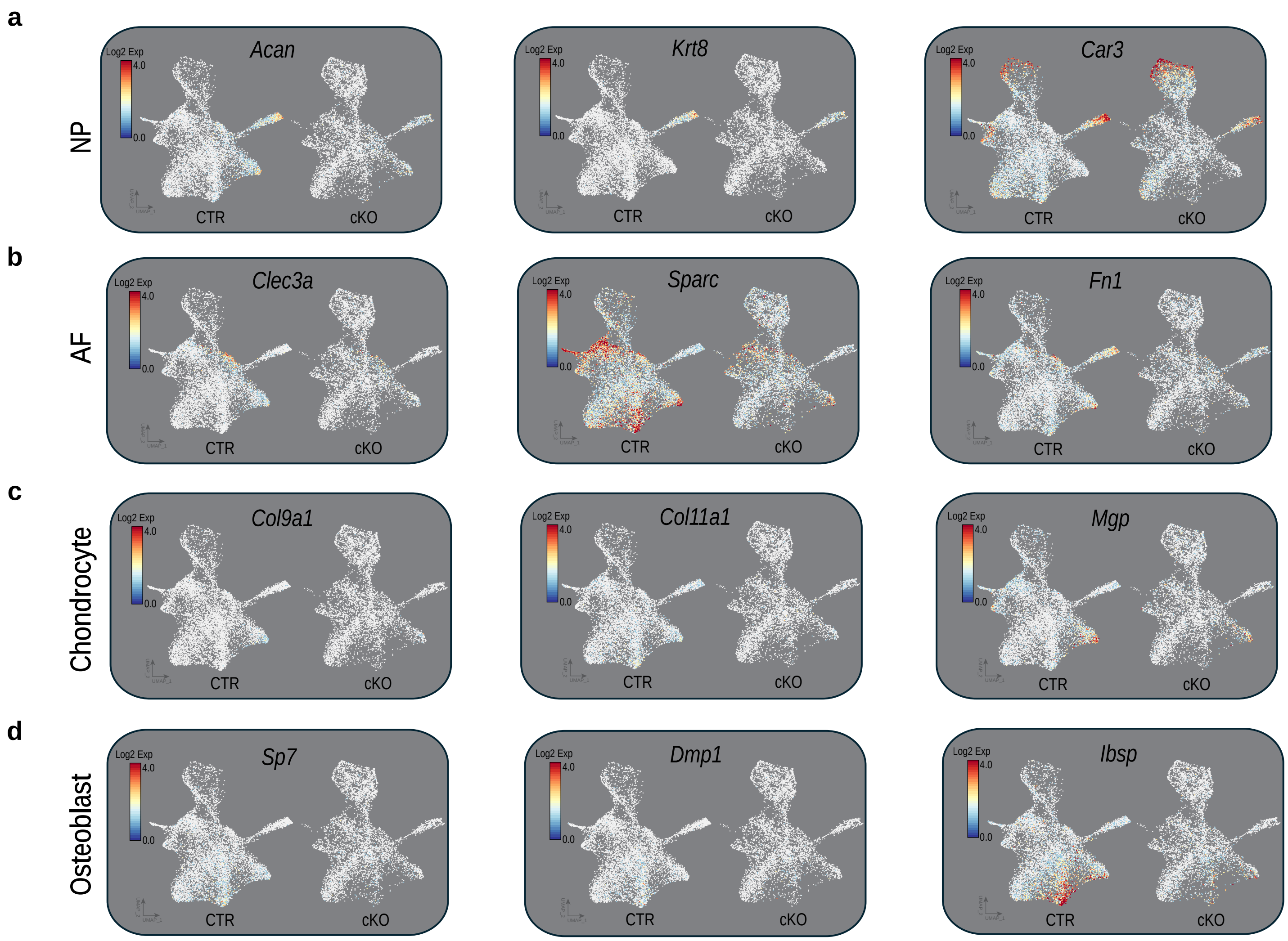

Supplementary Figure 3

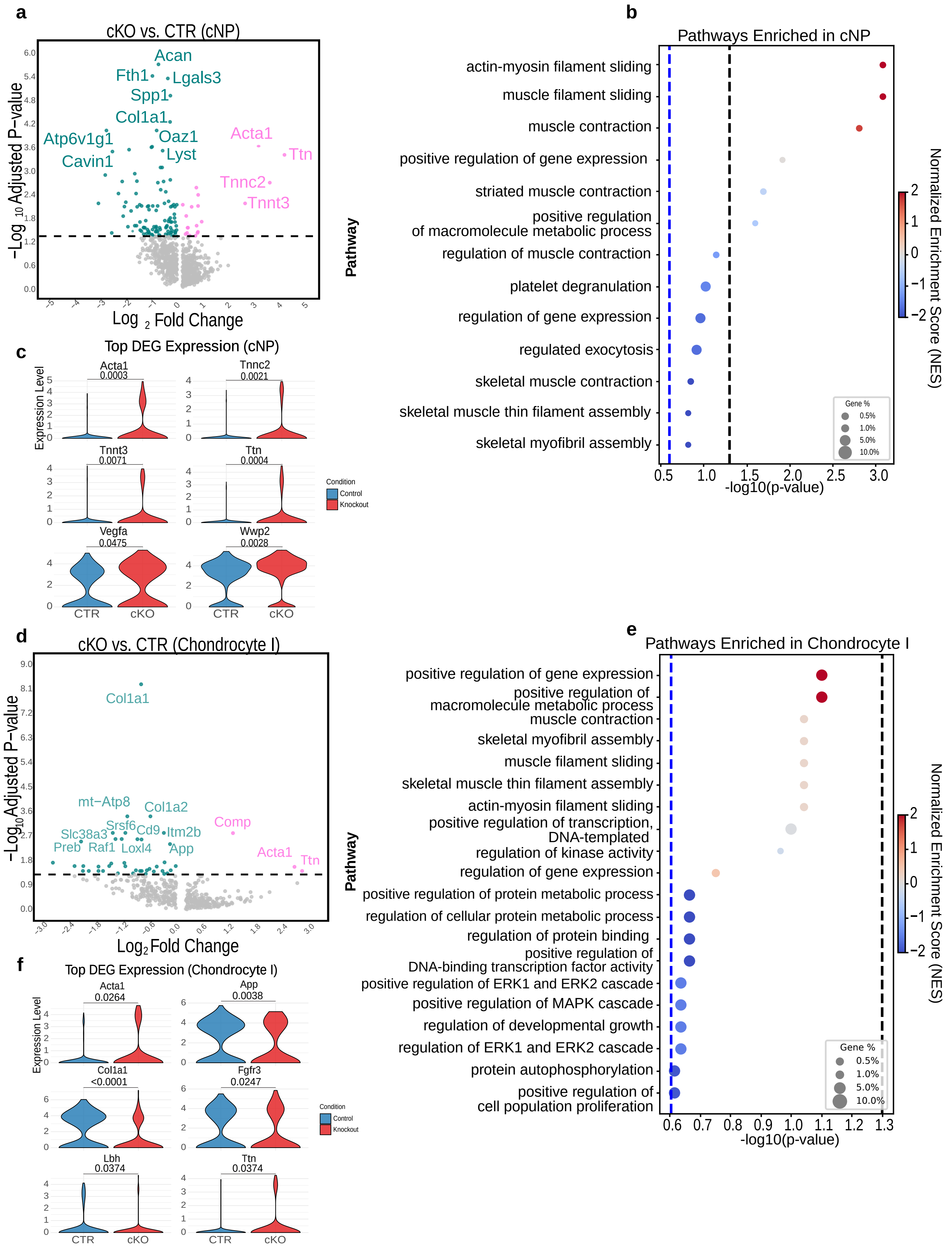

Supplementary Figure 4

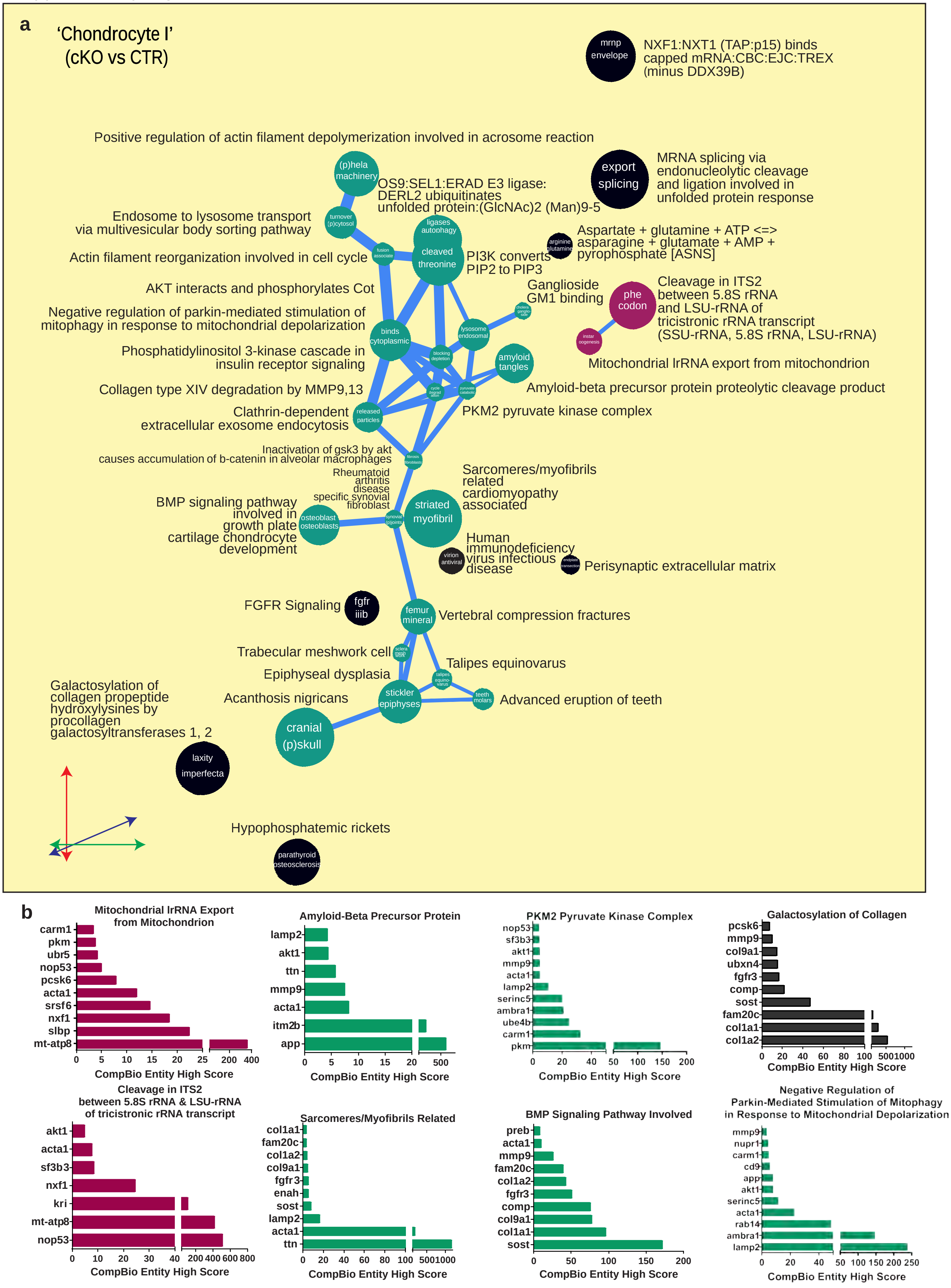

Supplementary Figure 5

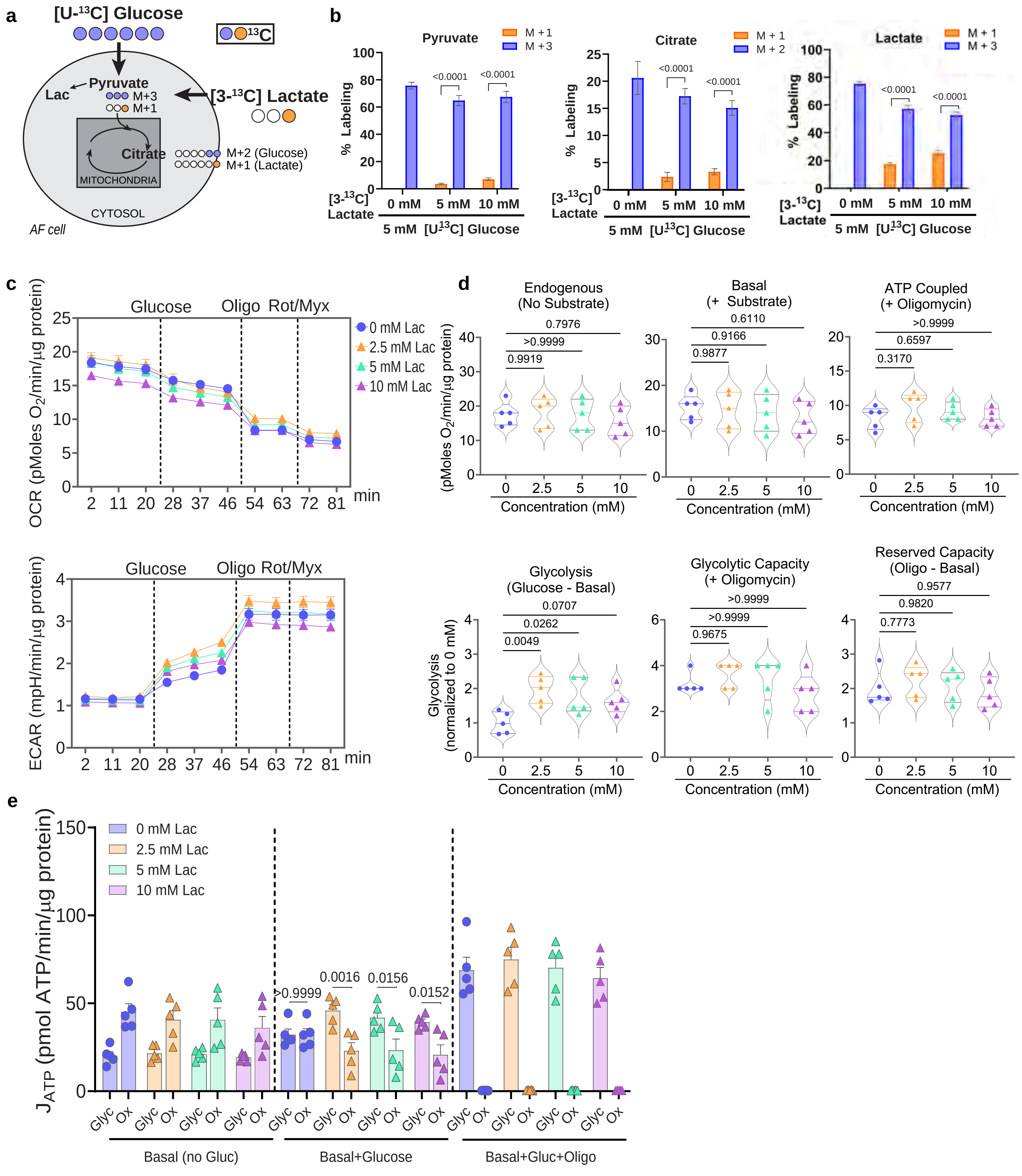

Supplementary Figure 6

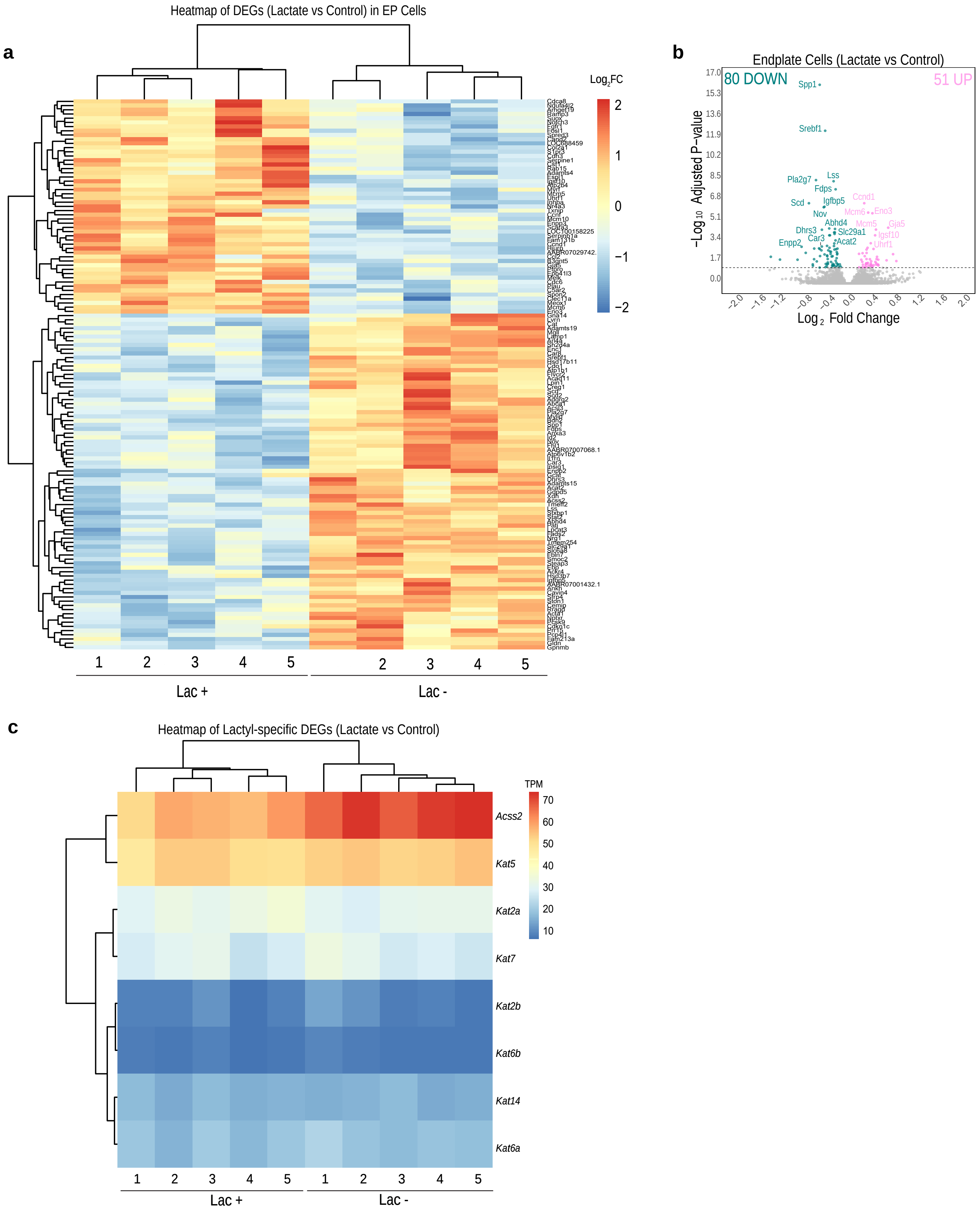

Supplementary Figure 7

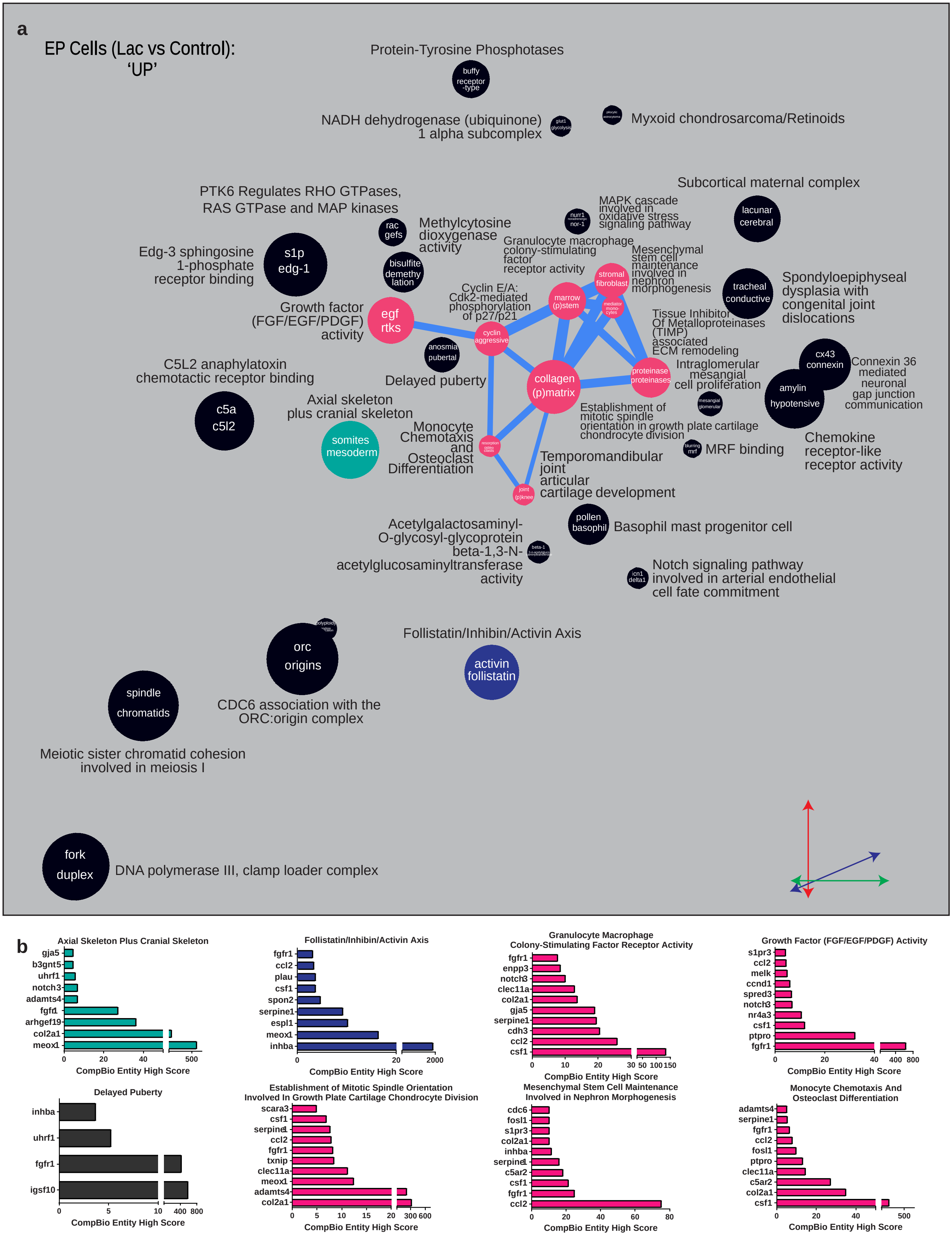
